# Survivor bias drives overestimation of stability in reconstructed ancestral proteins

**DOI:** 10.1101/2022.11.23.517659

**Authors:** Adam Thomas, Benjamin D. Evans, Mark van der Giezen, Nicholas J. Harmer

**Affiliations:** Living Systems Institute, Stocker Road, Exeter EX4 4QD, U.K.; Department of Biosciences, University of Exeter, Stocker Road, Exeter EX4 4QD, U.K.; Centre for Biomedical Modelling and Analysis, Stocker Road, Exeter EX4 4QD, U.K.; Department of Chemistry, Bioscience and Environmental Engineering, University of Stavanger, Richard Johnsens gate 4, 4021 Stavanger, Norway

**Keywords:** Evolution, survivor bias, modelling, protein evolution, protein engineering

## Abstract

Ancestral sequence reconstruction has been broadly employed over the past two decades to probe the evolutionary history of life. Many ancestral sequences are thermostable, supporting the “hot-start” hypothesis for life’s origin. Recent studies have observed thermostable ancient proteins that evolved in moderate temperatures. These effects were ascribed to “consensus bias”. Here, we propose that “survivor bias” provides a complementary rationalisation for ancestral protein stability in alignment-based methods. As thermodynamically unstable proteins will be selected against, ancestral or consensus sequences derived from extant sequences are selected from a dataset biased towards the more stabilising amino acids in each position. We thoroughly explore the presence of survivor bias using a highly parameterizable *in silico* model of protein evolution that tracks stability at the population, protein, and amino acid levels. We show that ancestors and consensus sequences derived from populations evolved under selective pressure for stability throughout their history are significantly biased toward thermostability. Our work proposes a complementary explanation of the origin of thermostability in the burgeoning engineering tools of ancestral sequence reconstruction and consensuses. It provides guidance for the thorough derivation of conclusions from future ancestral sequence reconstruction work.

## Introduction

Ancestral sequence reconstruction (ASR) traces changing protein sequences across evolutionary time (Akanuma 2017; Gumulya and Gillam 2017; Merkl and Sterner 2016; Scossa and Fernie 2021; Selberg, Gaucher, and Liberles 2021). Recently, ASR has been used to elucidate details about the evolution of several biochemical traits. These properties include substrate discrimination, specificity and plasticity (Babkova et al. 2017; Carter and Wills 2019; Pawlowski et al. 2018; Spence et al. 2021; Wheeler et al. 2018), trends in thermodynamic properties early in evolutionary time (Akanuma 2017; Akanuma et al. 2013; Butzin et al. 2013; Gaucher, Govindarajan, and Ganesh 2008; Hart et al. 2014; Hobbs et al. 2012; Okafor et al. 2018; Risso, Gavira, and Sanchez-Ruiz 2014; Schulte-Sasse et al. 2019), and the functional space within alternative evolutionary histories of protein families (Cole et al. 2013; Starr, Picton, and Thornton 2017). Other analyses have focused on structural characteristics of polypeptides, including the evolution of tertiary and quaternary protein structures (Hochberg and Thornton 2017; Lim and Marqusee 2018; Prinston et al. 2017; Schulte-Sasse et al. 2019), the evolution of complexity in multimeric proteins (Finnigan et al. 2012), and the evolution of viral capsid intermediate structures (Gullberg et al. 2010; Zinn et al. 2015). ASR has proved to be a useful tool for identifying proteins with novel properties for biocatalysis (Furukawa et al. 2020; Ma et al. 2022), vaccine design (Arenas 2020), directed evolution (Yu et al. 2022), and biologics (Knight et al. 2021), amongst other applications.

Thermostability of ancient proteins has been a particular ASR focus. ASR reconstructions of the most ancient sequences from protein families conserved across all kingdoms of life have consistently produced thermostable molecules (e.g. EF-Tu; (Butzin et al. 2013; Gaucher, Govindarajan, and Ganesh 2008; Hart et al. 2014; Okafor et al. 2018). Such proteins demonstrated a trend from high to low thermal stability from ancient life to modern life. These trends correlate with estimated historical terrestrial temperatures (Butzin et al. 2013; Gaucher, Govindarajan, and Ganesh 2008; Hart et al. 2014; Okafor et al. 2018). Many studies have concluded that early organisms inhabited a warmer Earth (Akanuma et al. 2013; Butzin et al. 2013; Gaucher, Govindarajan, and Ganesh 2008; Hart et al. 2014). This required proteins to fold and function under high temperatures. However, recent studies have uncovered thermostable proteins from lineages unlikely to have encountered high environmental temperatures in their evolutionary life history (Gumulya et al. 2018; Nicoll et al. 2020; Thomas et al. 2019; Trudeau, Kaltenbach, and Tawfik 2016), and that ancient seas may not have been as warm as previously believed (Galili et al. 2019). This suggests that not all thermostable ancestors are derived from the same conditions.

We recently reconstructed ancient carboxylic acid reductases (CARs) from the *Mycobacteria* and *Nocardia* that exhibited up to 35 °C increases in stability over their extant counterparts (Thomas et al. 2019). These ancestor stabilities do not fit the trends established by other paleotemperature studies (Akanuma et al. 2013; Butzin et al. 2013; Gaucher, Govindarajan, and Ganesh 2008; Hart et al. 2014). There is no prevailing evidence that any ancestors of *Nocardia* and *Mycobacteria* were thermophiles. The available evidence supports this family evolving from a common ancestor in the late Phanerozoic eon (<500 myo), when the earth was warming from a colder “snowball earth” state (Harland 1964; Lewin et al. 2016). Trudeau *et al*., 2016 reported a similar pattern with the serum paraoxonases (PON), whose ancestor was found to be up to 30 °C more temperature resilient than their modern-day counterparts. Ancient PONs exhibited superior folding properties when expressed in *E. coli* (Trudeau, Kaltenbach, and Tawfik 2016). Similar increases in stability were achieved by Gumulya *et al*. in the reconstruction of CYP3 cytochrome P450 mono-oxygenases (Gumulya et al. 2018). Both PONs and CYP3 are post-Cambrian innovations of Mammalia and Vertebrata respectively. There exists no evidence that any mammalian or vertebrate ancestor thermoregulated at the temperatures suggested by PON and CYP3 ancestor stabilities (Mackness and Mackness 2015).

To explain such stabilising effects, Gumulya *et al*. posited that vertebrate ancestors of CYP3 evolved in a warmer ocean environment. They propose that proteins subsequently approached mesophily by drift within recent evolutionary timescales (Gumulya et al. 2018). Trudeau *et al*. suggested that whilst environmental stresses or cellular defects in translation and chaperones might be responsible, the most likely explanation is that their ancestral PONs showed a strong consensus effect, especially in divergent surface residues (Trudeau, Kaltenbach, and Tawfik 2016). We also found that ancestral CARs exhibit bias towards the consensus sequence (Thomas et al. 2019). Whilst for well-conserved core residues, the consensus likely represents the ancestral sequence (Risso, Gavira, and Sanchez-Ruiz 2014), this is not necessarily the case for divergent positions (Trudeau, Kaltenbach, and Tawfik 2016). Consensus sequences are a proven sequence-driven method to engineer stabilising properties into enzyme families (Jia, Jain, and Sun 2021; Jones et al. 2020), hence the method has a stabilising effect on ancestral proteins (Durani and Magliery 2013; Kiss et al. 2009; Okafor et al. 2018; Sternke, Tripp, and Barrick 2019). Current explanations for the thermostable properties of consensus sequences assume that common amino acids at a position contribute to thermodynamic fitness more than other possible amino acids at that position (Porebski and Buckle 2016; Sternke, Tripp, and Barrick 2019; Ye, Yang, and Yu 2018). A comparison of ASR methods based on computational evolution (Williams et al. 2006) highlighted that the more popular maximum likelihood based methods have a tendency to produce ancestors more stable than the “true” ancestor, but suggested small stability increases of around 1.5 kJ mol^-1^, equivalent to a stability change of around 6 °C (Jaenicke 1999; Williams et al. 2006).

The proposed origins of stability in ancestral proteins apparently evolved from a mesophilic ancestor are therefore counterintuitive, incomplete, and insufficient for describing the underlying forces driving stabilization. It cannot be excluded that proteins with a recent origin evolved in warmer environments (Gumulya et al. 2018). This explanation becomes less parsimonious than an ASR-derived effect with every discovery of a new stable ancestor from mesophilic origins. We therefore explored an alternative hypothesis that there exists a “survivor bias”, which explains the stabilization of ancestral sequences in the absence of a stable ancestor. This has previously been explored to explain the stability of consensus sequences (Godoy-Ruiz et al. 2006). Briefly, the survivor bias hypothesis (Box 1) states that natural proteins incur a considerable fitness cost if their maximum folding temperature is below that of their immediate environment. As present-day proteins typically display stabilities (Δ*G*_f_) of -2 to -10 kJ mol^-1^ (average -5 kJ mol^-1^) (Bigman and Levy 2020; Godoy-Ruiz et al. 2006; Zeldovich, Chen, and Shakhnovich 2007), significantly destabilising mutations are selected against. Such mutations are therefore underrepresented in extant protein datasets, and stabilising residues are over-represented. Most ASR methods model evolution using mutation likelihoods that have been empirically derived from observed mutation patterns. These likelihoods are the global average of different distributions for each amino acid position in a protein. Survivor bias results in extant distributions of amino acids that over-represent the more stable amino acids in each position. This can confound some selections of the most likely amino acid at a node towards more stable options. Consequently, stabilising residues are over-represented in both consensus and ASR derived sequences.

### Box 1

**Overview of the survivor bias hypothesis**.

The survivor bias hypothesis is based on the following logical argument:

1. *The majority of amino acids are destabilising at any given site in a protein* (Taverna and Goldstein 2002). The distribution of effects on Δ*G*_f_ has been shown to be the sum of two Gaussians, with both having a destabilising mean (Faure and Koonin 2015; Tokuriki et al. 2007). A protein that has a natively folded structure has far more stabilising residues than the global distribution, with amino acids in any position expected to be populated according to their relative stability (Godoy-Ruiz et al. 2004; Godoy-Ruiz et al. 2005).
2. *The majority of proteins are marginally stable* (Bershtein et al. 2006; Goldstein 2011; Williams, Pollock, and Goldstein 2006). More stable proteins are possible and do indeed exist. However, as most mutations reduce stability, proteins tend to evolve a sequence that is “marginally” stable. The selective pressure occurs when further reductions in stability would render the protein unstable (Bershtein et al. 2006; Tokuriki and Tawfik 2009). The overall destabilising mean of all possible amino acids in each position and selective pressure to maintain folding act as counteracting forces.
3. *Sequence space for contemporary proteins over-represents stabilising or less destabilising amino acids compared to the global distribution*. Proteins must be at least marginally stable to survive natural selection. Contemporary proteins will therefore show a sequence distribution that over-samples amino acids in each position that are either stabilising or only mildly destabilising.
4. *Reconstructions based on contemporary proteins lack information as sequences selected against are absent*. Reconstructions generally assume that the sequences included are representative of all evolutionary paths that the ancestor could have taken. The full evolutionary web from an ancestral sequence includes many sequences that are unstable. These are selected against and are absent from our record. Reconstructions can consequently over-sample stabilising amino acids in each position. This has the potential to generate stable ancestral proteins when no stable ancestor is predicted to have existed (Gumulya et al. 2018; Lewin et al. 2016; Nicoll et al. 2020; Thomas et al. 2019).

To test the survivor bias hypothesis, we developed an *in silico* model of sequence evolution called “PESST” (Protein Evolution Simulations with Stability Tracking)^1^. PESST evolves a population of protein sequences and tracks the changing stability of these sequences. PESST was designed as a sequence evolver that follows standard amino acid evolution, generates phylogenies *de novo*, and focuses on the integration of environmental constraints on the evolving population fitness. By observing the outcomes of simulated evolution, we identified that simultaneous effects from both the destabilising force of drift and the stabilising force of a stability threshold are driving bias in ASR. We observed that simulated ancestors showed large increases in stability, far higher than any stability sampled throughout their evolutionary history. There is a significant correlation of stability with ancestor “age”. The simulated populations produced consensus proteins that were different in sequence from ancestors, and significantly stabilized. PESST provides a sandbox to test evolutionary hypotheses. It provides strong support that survivor bias can cause protein stabilization in sequence alignment-driven protein engineering tools unless caution is taken. These data suggest that ASR starting from diverse populations is a powerful engineering tool for the biasing of sequences towards stability, irrespective of a protein’s evolutionary history.

## Results and Discussion

### The PESST algorithm tracks stability as the protein evolves

We firstly checked that our model (PESST v1.1) tracks expected population stability over time. We confirmed that the mean free energy of a population of sequences without any selection pressure evolves to the equilibrium (*ϵ*) of the stability space. Example simulations were performed where the starting sequences were selected to be of very low stability (Figure 1A), of average stability (Figure 1C), or highly stable (Figure 1E). We simulated five repeats of each scenario. In all cases, the mean population stability converged on the mean stability of all available protein sequences within 5 000 simulated generations with a mutation rate of 0.002 bases/generation. The convergence towards the mean occurs more rapidly from starting sequences that are highly stable than those with very low stability (Figures 1A, E). This is consistent with the expected energy distribution being skewed towards destabilising amino acids.

**Figure 1:**
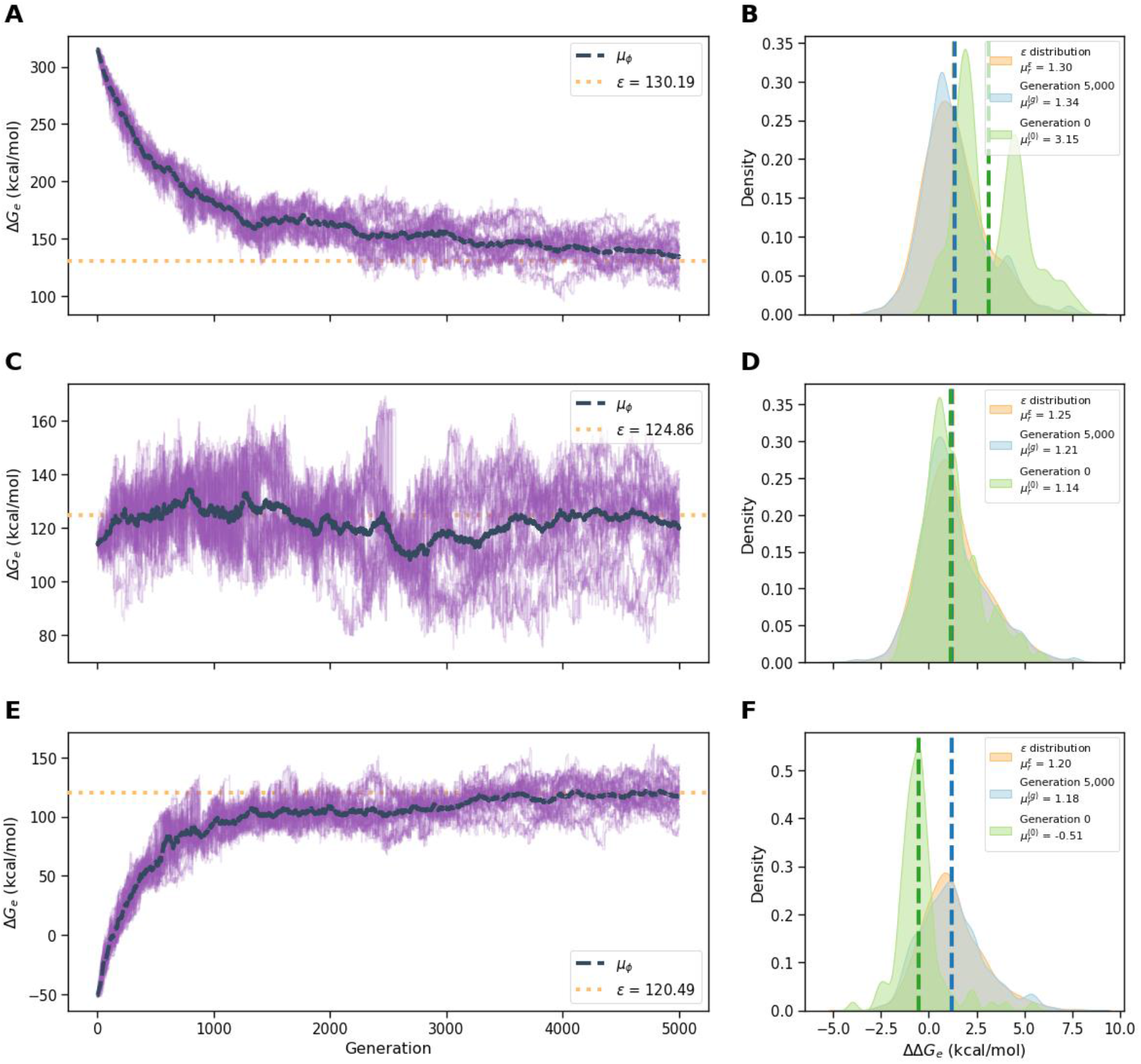
Mean stability of the population simulated in PESST tends toward *ϵ* during evolution. Representative stability traces for PESST simulations over 5 000 generations. Δ*G*_0_ values were modelled for each amino acid following Tokihuri et al. (2007). The mutation rate (*p*_*m*_) was set to 0.002 (mutations per position per generation). Simulations were initialised with protein sequences chosen from the three amino acids with the highest locus ΔΔ*G*_0_ at each position (**A**,**B**), the sixteen amino acids closest to the mean locus ΔΔ*G*_0_ at each position (**C**,**D**), or the three amino acids with the lowest Δ*G*_0_ at each position (**E**,**F**). In the left-hand graphs, each one of 52 clones in the dataset was tracked independently and simultaneously by PESST. The dashed black bold line represents the mean Δ*G*_0_ of the population. The orange horizontal dashed line represents the predicted value of ***ϵ***, the global mean sum of Δ*G*_0_ across all amino acids. The right-hand graphs show the distribution of locus ΔΔ*G*_0_ values. The starting and ending distributions for the population are shown in green and blue respectively, whilst the distribution of all possible amino acids in all positions is shown in orange. The results show that the stability of PESST simulated protein populations approaches the predicted value of *ϵ* in all three of these settings.

To confirm that these means were capturing the expected variation in Δ*G*_0_ values across the population, we tracked the distribution of ΔΔ*G*_0_ values per amino acid position (“locus ΔΔ*G*_0_”) in the evolving dataset (Figure 1B, D, F). As the simulation reaches the observed equilibrium, it is expected that the distribution of locus ΔΔ*G*_0_ in the simulated population will tend to reflect the initially defined matrix of locus ΔΔ*G*_0_ values. We observed that the population mean of ΔΔ*G*_0_ values 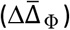 rapidly converged to the initial matrix mean Δ*G*_0_ (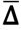 Figure 1). The evolving locus ΔΔ*G*_0_ distribution converges to the initial matrix distribution, but more slowly, in the three scenarios tested (Figure 1B, D, F). After this convergence there is no statistically significant difference between the two distributions (*p* > 0.3 by generation 5 000 in all tested cases; Kolmogorov-Smirnov test).

### A stability threshold biases the distribution of locus Δ*G*_0_ values in the evolving dataset

Our survivor bias hypothesis proposes that the overall protein stability will tend towards a global Δ*G* (Δ*G*_0_) of zero. At this point protein folding becomes unfavourable, imposing a fitness cost on the host organism (Harms and Thornton 2013; Khersonsky et al. 2018; Williams, Pollock, and Goldstein 2006). We tested this using PESST by selecting against clones whose global stability becomes unfavourable (Δ*G*_e_ > 0). We simulated evolution starting from a highly stable initial sequence (Figure 2). The population evolved as before for the first 250–500 generations. Once unstable clones arise and are eliminated from the population, the population mean stability tends towards an equilibrium (Figure 2B). This equilibrium has a range of -1 to -8 kcal mol^-1^, with most values between -2 and -3 kcal mol^-1^.

**Figure 2:**
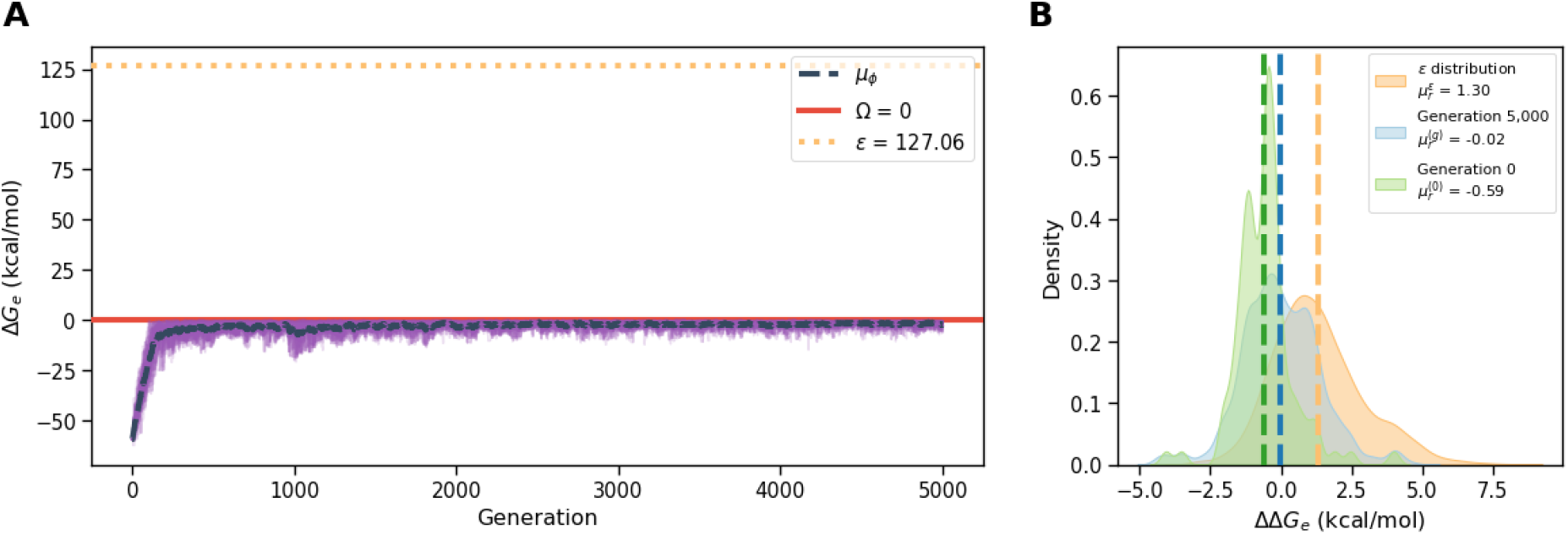
Selection against unstable clones leads to an equilibration mean protein stability with a positive bias in locus ΔΔ*G*_0_ distribution. Data represents the simulation of 5 000 generations, with a stability threshold of Δ*G*_0_ = 0 imposed. Clones that exceed the threshold are removed and replaced *in populus*. Five simulations were initiated with highly stable starting sequences. A representative simulation is shown, in the same fashion as Figure 1. **A:** Each clone in the dataset was tracked independently. The tight dashed black bold line represents the mean Δ*G*_0_ of the population. The orange horizontal dashed line represents the predicted value of ***ϵ***, the global mean sum of Δ*G*_0_ across all amino acids. Average stability across the population is maintained at equilibrium at ∼ Δ*G*_e_ = -2 kcal mol^-1^. **B:** Distribution of locus ΔΔ*G*_0_ values. The starting and ending distributions for the population are shown in green and blue respectively, whilst the distribution of all possible amino acids in all positions is shown in orange. At generation 5 000, the extant amino acid probability matrix 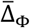 does not converge to the initial global matrix 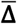 (*p* << 0.0005)^2^. Negative locus ΔΔ*G*_0_ values are significantly overrepresented, and destabilizing Δ*G*_0_ values significantly underrepresented (Supplementary Figure 2).

This equilibrium is maintained by bidirectional pressures. Neutral evolution tends to reduce the mean stability due to the destabilising initial matrix **Δ**. Selection against clones that become unstable provides a stabilising pressure. These observations confirm that PESST successfully simulated the expected population behaviour in the context of a biologically relevant stability threshold. This is consistent with previously reported observations of protein evolution that underpin our survivor bias hypothesis (Faure and Koonin 2015; Goldstein 2011; Pucci and Rooman 2016; Tokuriki and Tawfik 2009).

We propose that when the stability criterion is imposed, the bidirectional pressure on protein stability will cause an over-representation of amino acids with a strongly negative locus ΔΔ*G*_0_ in the evolving population. Our simulations demonstrate that at equilibrium, the extant amino acid probability matrix 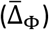 does not converge to the initial global matrix (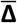 Figure 2B). The expected over-representation of stabilising amino acids is observed. Across our replicates, we observed that the most destabilising amino acids titrate out of the population. This is likely due to the large stability penalty that these impose (Figure 2B). These most destabilising residues impose a penalty in the range 3–8 kcal mol^-1^, sufficient to cause an average clone at equilibrium to be removed from the population. This is consistent with previous analyses of factors underlying marginal stability in proteins (Goldstein 2011; Tokuriki and Tawfik 2009). Locus ΔΔ*G*_0_ distributions with the stability criterion imposed over-represent stabilising locus ΔΔ*G*_0_ values in comparison to simulations without this criterion (*p* << 0.0001; Supplementary Figure 1). This limitation on the evolutionary space that the protein can sample is consistent with Goldstein’s study, which showed that rare stabilising mutations tend to be fixed in positions where there is no fitness benefit to amino acids that provide less stability (Goldstein 2011). A similar shift in locus Δ*G*_0_ distribution compared to the global distribution was observed for mutations in extant proteins (Tokuriki et al. 2007). This work provides further evidence that the bidirectional evolutionary pressures on an evolving protein induce stability biases into the locus ΔΔ*G*_0_ distribution even at equilibrium.

### Ancestral reconstruction and consensus sequences overestimate protein stabilities when deriving sequences from evolutionary simulations with a fitness threshold imposed

We propose that the effect of limiting the evolutionary space that a protein sequence can sample (i.e. a stability criterion) will cause ancestral protein reconstruction to bias towards stability. This may result in an “ancestral” protein more stable than any sampled in the evolutionary history of the protein family. To test this hypothesis, we reconstructed “ancestors” of PESST simulated evolution using the commonly used CodeML reconstruction algorithm in PAML (Yang 2007). Our null hypothesis (no induced bias) is that ancestors will share the stability space of the dataset’s evolutionary history. We simulated evolution using the locus Δ*G*_0_ distribution identified by Tokihuri et al. (Tokuriki et al. 2007) (**Experiment 4a**); using a distribution with the same shape but the mean Δ*G*_0_ halfway between this and zero (**Expt 4b**); a distribution with the same spread and mean Δ*G*_0_ equal to zero (**Expt 4c**); and the inverse of Expt 4b (i.e., a stabilising distribution; **Expt 4d**). These alternative distributions allow us to test the strength of the effect. The starting sequences were selected to be marginally stable to avoid introducing bias into the history.

Where the global Δ*G*_0_ distribution is destabilising, the evolutionary histories converge to an equilibrium between -1 to -2 kcal mol^-1^ (Expt 4a) or -4 to -6 kcal mol^-1^ (Expt 4b) over 2 000 generations. As we previously observed, the extant amino acid probability matrix 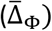 does not converge to the initial global matrix 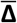. The statistical significance of the difference between the distributions is greater in Expt 4a (*p* = 10^−110^ to 10^−186^) compared to Expt 4b (*p* = 10^−48^ to 10^−57^). With a global distribution centred on Δ*G*_0_ = 0 (Expt 4c), an equilibrium was not reliably maintained, and a broad stability space is sampled (Supplementary Figure 2A). In these conditions the bidirectional selection pressures are both weak. With an artificial stability distribution centred on stabilising Δ*G*_0_ values, a broad sequence space is sampled (Supplementary Figure 2B). For both experiments 4c and 4d, the evolved locus Δ*G*_0_ distribution is the same as the global Δ*G*_0_ distribution (*p* > 0.999). For Expt 4a, we investigated the ratio between stabilising and destabilising mutations. Surprisingly, once an equilibrium has been achieved (∼250 generations), stabilising mutations were as common as destabilising mutations (Supplementary Figure 3). We could not observe mutations that caused sequences to become unstable and so selected against; and expect that our observed ratio reflects the purifying effect of selection against these mutations.

Sequence alignments at generation 2 000 had 30–50% pairwise sequence identity. Phylogenies generated from these alignments (Guindon et al. 2010) were well supported for most nodes (Supplementary Figure 4; Supplementary files). We reconstructed ancestors from these phylogenies (Yang 2007). “Ancestor” stabilities were calculated from the global Δ*G*_0_ matrix for each simulation. We measured the impact of the bidirectional selective pressures by taking the sum of folding energies (Δ*G*_f_) for the 52 extant clones and 51 ancestors and assessing the proportion of this population stability attributed to the extant and ancestor clones. Our null hypothesis is that the extant and ancestor proteins will contribute equally to the population Δ*G*_f_. This outcome would suggest that the stability criterion does not affect ancestral reconstruction. For the simulations where the initial global matrix 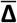 has a positive Δ*G*_0_, the normalized stability space is dominated by the ancestral clones (78% for Expt 4a; 61% for Expt 4b; Figure 3A). In contrast, where the initial global matrix averages to zero (Expt 4c) or is stabilising (Expt 4d), there is little effect (53% and 50% respectively). Previous simulations of protein evolution similarly observed that reconstructions of evolving proteins have overestimated stability (Williams et al. 2006; Williams, Pollock, and Goldstein 2006).

**Figure 3.**
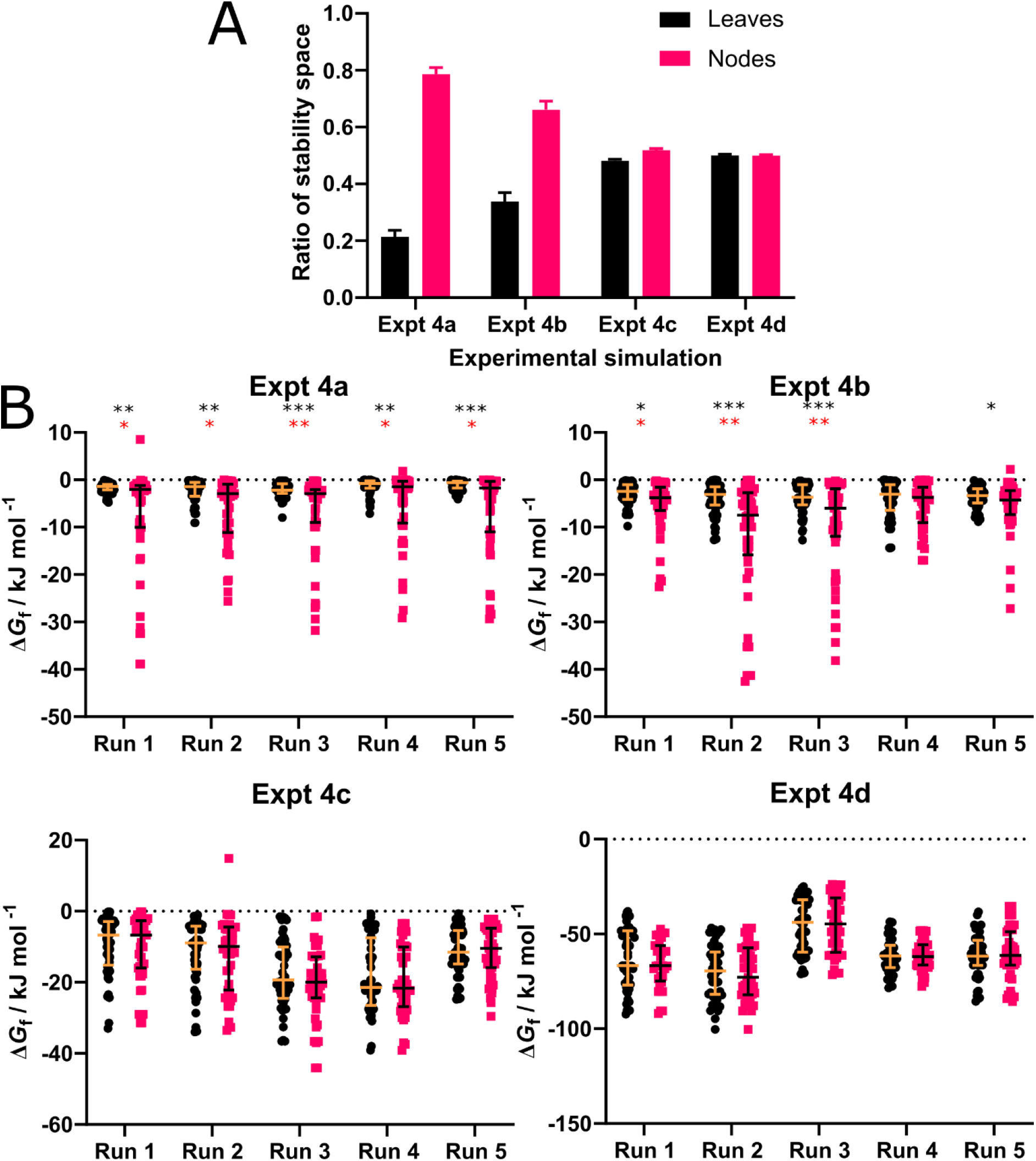
Survivor bias drives increasing stability in ancestral proteins. Figures represent analyses of stability space across ancestral and extant sequences. Each simulation was repeated five times. Evolution was simulated using experiment locus ΔΔ*G*_0_ distributions (Tokuriki et al. 2007) (**Expt 4a**); this distribution with the mean Δ*G*_0_ halfway between the experimental values and zero (**Expt 4b**); this distribution with mean Δ*G*_0_ equal to zero (**Expt 4c**); and the inverse distribution to Expt 4b (i.e., a stabilising distribution; **Expt 4d**). Each simulation evolved for 2 000 generations. **A**: Global stability space is biased toward nodes when the global stability distribution is destabilising. Tree stability space for each run was normalized and proportioned between node (ancestor; pink) and leaf (extant; black) stability. Mean proportion values over replicates shown. Error bars represent standard error of the mean. **B**: Columnar scatter plots comparing the distribution of leaf stabilities and node stabilities, When the average locus Δ*G*_0_ is negative, node stabilities are overestimated. Comparisons of the distributions were undertaken on a run-by-run bases with both Welch’s *t*-test (black asterisks) and Mann-Whitney U test (red asterisks). Black horizontal bars represent the mean and range of the datasets. Asterisks represent the degrees of significance (*** = *p* < 0.0005; ** = *p* < 0.005; * = *p* < 0.05).

To further understand the observed stability bias, we compared the distribution of stabilities in extant and ancestor populations on a per-simulation basis. There is a clear disparity between the Δ*G*_f_ values sampled in the two populations when experimentally determined locus ΔΔ*G*_0_ values are used (Figure 3B, Expt 4a). Ancestors (nodes) sample a range of Δ*G*_f_ values between 2.8 and 10.2 times wider than extant clones (leaves). All simulations show a highly significant difference between nodes and leaves (*p* = 6×10^−4^ to 5×10^−5^; Supplementary table 5). In each simulation, the most stable ancestors show stabilities much higher than any sampled in the evolutionary history of their parent population (Supplementary Figure 5; supplementary files). The stabilization observed (5–25 kJ mol^-1^) is equivalent to an increase in temperature stability of 20–100 °C, depending on the precise protein structure (Jaenicke 1999; Vicatos, Roca, and Warshel 2009; Williams et al. 2006). This is greater than the stabilising effect previously observed (Williams et al. 2006). However, this study evolved proteins only to the equilibrium point whilst our evolution extends for many generations beyond this. This highlights that reconstructions deeper into the past can show greater effects. Artificially reducing the average fitness penalty of locus Δ*G*_0_ by half significantly reduced this effect (Figure 3B, Expt 4b), with only four of the five simulations showing a significant difference (*p* = 0.15 to 3×10^−5^). When the average locus ΔΔ*G*_0_ values were zero (Expt 4c) or negative (Expt 4d), there was no difference between leaves and nodes (Figure 4B; *p* > 0.1 in all cases). In these cases, ancestral stabilities are not overestimated. These data lead us to reject our null hypothesis as the ancestors demonstrate stabilities that are not sampled by the population in their evolutionary history. This effect requires the bidirectional pressure of a requirement for global protein stability and the average locus ΔΔ*G*_0_ being destabilised.

**Figure 4:**
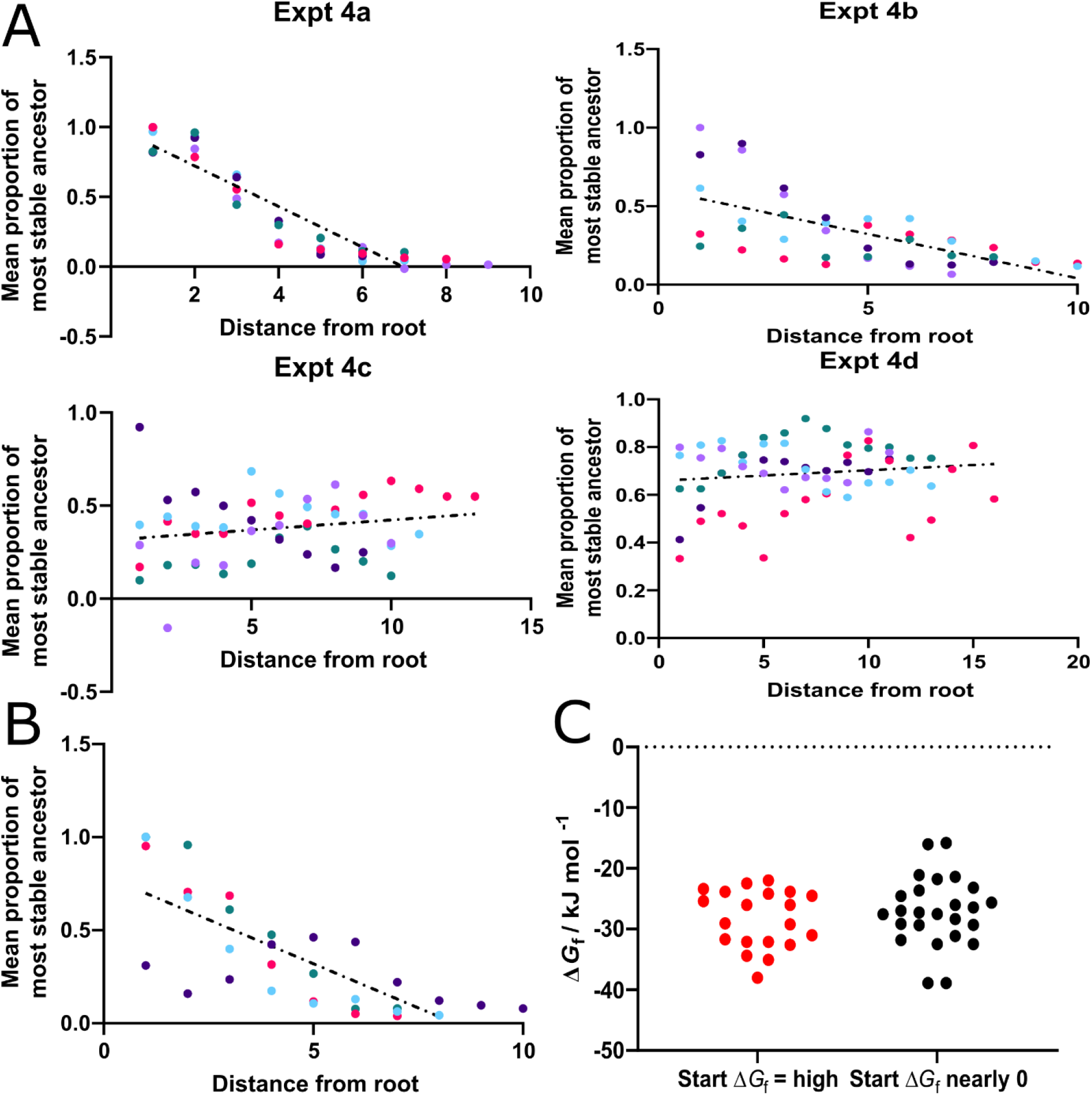
Survivor bias manifests as older ancestors showing greater stability. Data derived from the experiments outlined in Figure 4. **A**: Scatter plots of normalized stability with respect to distance from the root of the tree. Node distance from the root was calculated from cladograms output from PAML as the number of nodes preceding the node of interest. Node stabilities were normalised to 1 (most stable node). Each of the five replicate simulations is individually coloured. A strong negative correlation is seen when the mean locus Δ*G*_0_ is destabilising (Expt 4a, 4b), but not when this is neutral (Expt 4c) or stabilised (Expt 4d). **B**: Scatter plot analysed as above for a simulation of protein evolution initiated with a highly stable protein. **C**: Grouped scatter of the five most stabilising Δ*G*_f_ values from each of five simulations starting at high stability (red), or with a marginally stable starting sequence (black). Mann Whitney U test determined that the populations were not significantly different.

Many past ASR studies have shown that sequences trend toward increased stability as they are reconstructed from more ancient nodes (Akanuma et al. 2013; Butzin et al. 2013; Gaucher, Govindarajan, and Ganesh 2008; Hart et al. 2014; Hobbs et al. 2012; Okafor et al. 2018; Risso, Gavira, and Sanchez-Ruiz 2014). We analysed the effect of distance from the root of our simulated reconstructions on ancestor stability (Figure 4A). Significant negative correlations (*p* < 0.0001) were observed for Experiments 4a and 4b (*r* = -0.91 and -0.65 respectively). In contrast, when the mean locus ΔΔ*G*_0_ value was not destabilising, no significant correlation was observed (*p* > 0.1 for Experiments 4c and 4d; Figure 4A). These simulations demonstrate that evolution under these bidirectional pressures is sufficient to produce ancestral reconstructions where stability correlates with age. This raises the possibility that in some cases, the thermostability of ancient proteins is not indicative of their environmental temperature, but instead of survivor bias. We therefore urge caution when drawing such conclusions from ASR studies.

To further illustrate this need, we simulated evolution starting at high stability (Expt 4e; Supplementary Figure 6) and reconstructed the ancestors of these simulations. Correlations in ancestor stability over time were calculated as before. A significant negative correlation was observed (r = -0.76, p < 0.0001; Figure 4B). This effect was similar to that observed with a low starting stability (Figure 4A, Expt 4a). Comparing the five most stable ancestors in these two simulations showed no significant difference (Figure 4C; *p* = 0.42). Confidently distinguishing whether trends of increasing stability are caused by survivor bias or a true evolutionary history requires additional evidence about the evolutionary history of the protein in question (e.g. correlation with isotope ratios deposited in ocean cherts) (Akanuma 2017; Garcia et al. 2017; Gaucher, Govindarajan, and Ganesh 2008). Others have suggested that only tracking single proteins is an unreliable way to infer conclusions of the ancient biosphere (Hart et al. 2014). Methods and guidelines for performing such reconstruction experiments with apt rigor are outlined in (Kacar et al. 2017). It must be noted that these conclusions are based on a model and need confirming by experimental evolution to create novel, stable proteins.

An alternative alignment-based tool for engineering protein stability is to use a consensus protein (Durani and Magliery 2013; Kiss et al. 2009; Okafor et al. 2018). Consensus proteins take the most common amino acids in each position in extant proteins. On average, individual substitutions to the consensus are as likely to be stabilising as destabilising (Polizzi et al. 2006; Sullivan et al. 2012). Amino acids tend to populate alignments at rates proportional to their stability (Godoy-Ruiz et al. 2004; Godoy-Ruiz et al. 2005). The increased stability of global consensus sequences is ascribed to the sum of many small favourable interactions between consensus residues (Sternke, Tripp, and Barrick 2019). As we have shown here, survivor bias titrates out residues with a destabilising locus Δ*G*_0_. This should over-populate the pool of extant proteins with stabilising residues, biasing consensus sequences towards increased stabilities, as has been shown experimentally (Sternke, Tripp, and Barrick 2019). To test this hypothesis, we calculated consensus sequences for generation 2 000 for experiments 4a–4d. Survivor bias was sufficient to generate consensus sequences with far greater stabilities than those sampled across the population’s evolution (*p* < 0.00001 for Expt 4a, 4b, 4c and 4e; Mann-Whitney U-test; Figure 5). When the global mean locus Δ*G*_0_ is stabilising (relieving the survivor bias effect), consensus sequences were slightly less stable than the extant population (*p* = 0.03). Consensus sequences showed significantly greater sequence similarity to the oldest ancestors than extant sequences when the survivor bias effect is in operation, but not when it is not (*p* = 3×10^−11^, 0.00005, 0.00004, and 0.9 for Expt 4a–4d respectively; Supplementary Figure 7). These data support the hypothesis that consensus sequences are stabilized by the introduction of ancestral residues, albeit in an evolutionary history dependent manner (Porebski and Buckle 2016; Ye, Yang, and Yu 2018).

**Figure 5:**
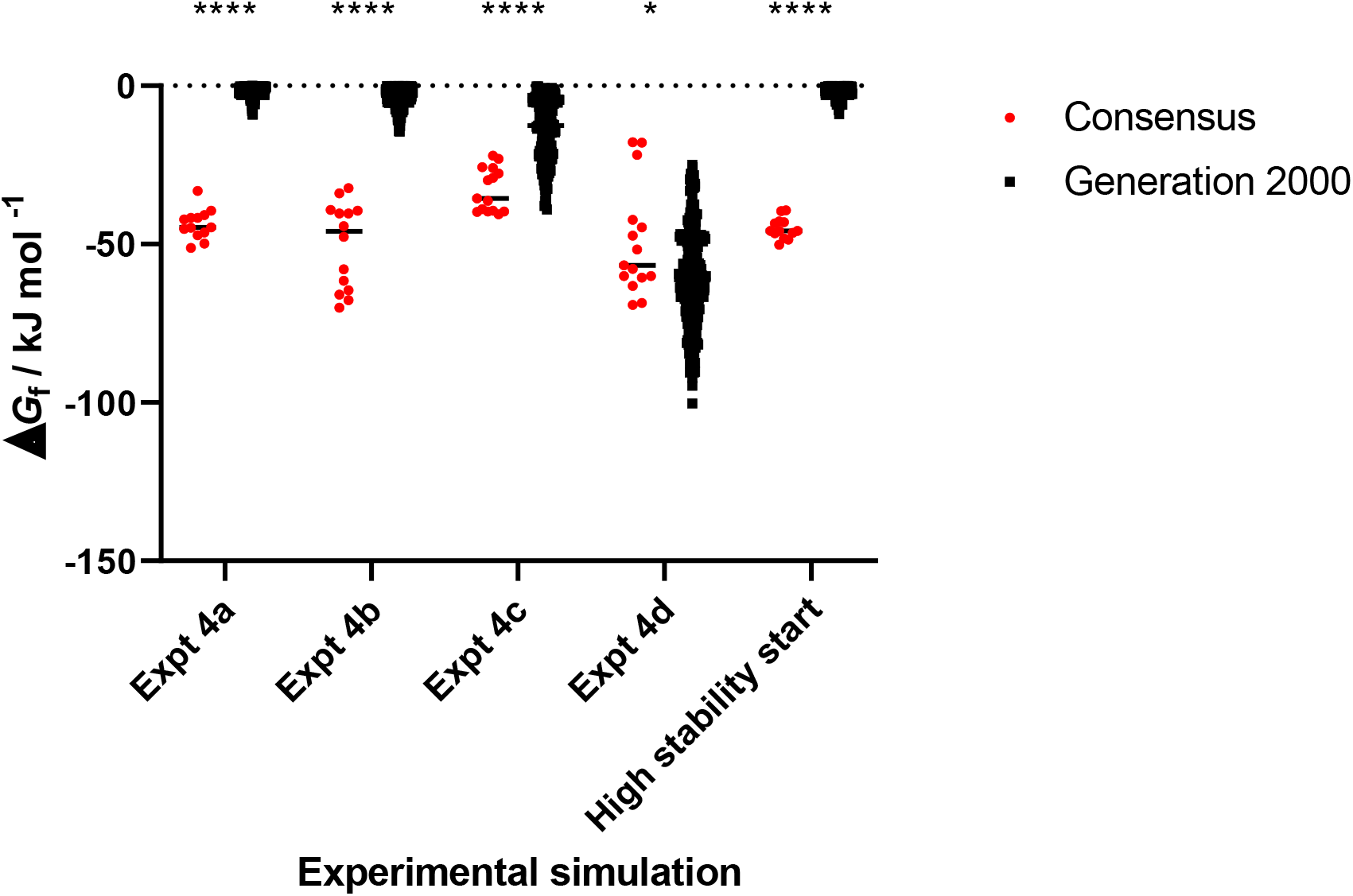
Survivor bias drives stabilization of consensus sequences. Grouped scatter plots show the stability of consensus (red) sequences derived from generation 2,000 (black) of **Experiments 4a–e** (Figures 4 and 5). Either two or three consensus sequences were determined for each simulation by PESST depending on the strength of the consensus. The stability of the consensus and generation 2 000 sequences were compared using the Mann-Whitney *U*-test. * *p* < 0.05; **** *p* < 0.00001.

These data suggest that all consensus sequences from real data are likely to show a bias to stability. Stabilising residues are expected to be over-represented in extant protein families as they are subject to survivor bias, as most locus ΔΔ*G*_0_ values are negative (Tokuriki et al. 2007). Selecting residues from alignments of extant proteins therefore distorts these probabilities by offering a biased selection space, especially in divergent positions (Trudeau, Kaltenbach, and Tawfik 2016). A recent study of consensus engineering found that 75% of families tested show higher stability than any extant counterpart (Sternke, Tripp, and Barrick 2019). As both consensus and ASR methods manifest stabilisation due to survivor bias unless this is accounted for, ASR-based stabilisation should also be robust for a broad range of protein families (Clifton et al. 2017; Wilding et al. 2017; Zakas et al. 2017). Recent examples include the PON, CYP3, FMO, and CAR families (Gumulya et al. 2018; Nicoll et al. 2020; Thomas et al. 2019; Trudeau, Kaltenbach, and Tawfik 2016). Furthermore, “young” ancestors may show stabilisation properties more reflective of more basal ancestors. This may add complexity to future ASR studies probing relationships between evolutionary history and stability of ancient proteins, especially in more divergent sequence alignments.

## Conclusions and perspective

> *“…life is one big lottery in which only the winning numbers are visible*.*”*
>
> --- Jostein Gaarder, *Sophie’s World*

In this study, we provide strong evidence for the hypothesis that “survivor bias” can explain the (thermo)stability of ancestral and consensus proteins derived from extant sequences. Our novel algorithm (PESST) simulates evolution whilst tracking protein stability. Our simulations demonstrate the effect of the bidirectional pressures of most mutations being destabilising and need for a negative global protein free energy of folding. Reconstructions from a population evolving under these bidirectional pressures show strong, significant correlation between apparent ancestor “age” and stability. We find that consensus sequences are similarly stabilised, accessing the same stabilising bias through a different route. Our work explains how reconstructions of protein families without a high temperature evolutionary history still produce thermostable ancestors (Thomas et al. 2019; Trudeau, Kaltenbach, and Tawfik 2016). Our results provide a broader understanding of the basis for alignment-based stability engineering methods (Gumulya et al. 2018; Sternke, Tripp, and Barrick 2019; Trudeau, Kaltenbach, and Tawfik 2016). We provide further evidence that ASR and consensus methods have potential as ubiquitous engineering tools, enabling the engineering of stability regardless of a protein’s evolutionary history.

## Methods

### Model description

PESST is a protein evolution simulator (summarised in Algorithm 1 with default parameters in Table 1) which simulates a fixed population (Φ), of *N* proteins evolving over *G* generations. Each protein, *η*, consists of *R* amino acids (methionine followed by *R* − 1 randomly chosen amino acids from 20 candidates) with an associated thermal stability defined as: 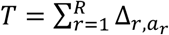, where Δ_*r,a*_ is the change in thermodynamic stability conferred by amino acid *a* at location *r* (i.e. “locus ΔΔ*G*_0_”; Supplementary Figure 8). The global set of Δ_*r,a*_ stability changes are randomly drawn from a distribution based on one or more Gaussians of defined mean, variance and proportion (Tokuriki et al. 2007), with an optional skew, ∼𝒩(*μ*_*n*_, *σ*^2^_*n*_, *skew*_*n*_, *P*_*n*_). For simplicity epistasis was not modelled since the majority of amino acid stability contributions are approximately additive (Bloom et al. 2005).

**Table 1:** Parameters in PESST. Note that for the studies reported here, a dual stability distribution was used following (Tokuriki et al. 2007).

| Parameter | Description | Typical Value | Notes |
| --- | --- | --- | --- |
| $G$ | Number of generations | 5 000 | $[1, \infty)$ |
| $T_0$ | Initial protein stability | (-5, -50) | {(lower, upper),<br>“low”: very unstable,<br>“mid”: average stability,<br>“marginal”: stability just<br>greater than threshold $\Omega$,<br>“high”: very stable} |
| $\Omega$ | Stability threshold | 0 | Stable := $T \leq \Omega$<br>Unstable := $T > \Omega$ |
| $\mu$ | Mean stability change ($\bar{\Delta}$) | 0.54 and 2.05 | $\mu \approx \frac{1}{R \cdot N} \cdot \sum_r \sum_a \Delta_{r,a}$ |
| $\sigma$ | Standard deviation of $\Delta$ | 0.98 and 1.91 | $\sigma \approx \text{var}(\Delta)^{\frac{1}{2}}$ |
| $k$ | Skew of $\Delta$ | 0 | |
| $N$ | Population size | 52 | |
| $n_{roots}$ | Number of roots | 4 | $2 < n_{roots} < N$ |
| $R$ | Protein length | 100 | $[1, \infty)$ |
| $p_{inv}$ | Proportion of invariant sites | 0.1 | $[0, 1)$ |
| $p_m$ | Probability of mutation | 0.002 | Total mutations := $p_m \cdot R \cdot N$ |
| $p_{death}$ | Probability of death | 0.05 | $[0, 1)$ |
| $\kappa$ | Gamma distribution shape | 1.90 | |
| $\theta$ | Gamma distribution scale | 0.53 | $1/\kappa$ |
| $\Gamma_i$ | Number of gamma iterations | 50 | |
| $\Gamma_s$ | Number of gamma samples | 10 000 | |

The population is initialised with proteins with a defined stability (“High”, “Low”, “Medium”, or within a desired range). Proteins evolve according to a simple uniform clock, with a fixed probability of mutation (*p*_*m*_) for each amino acid in every generation, except for a defined proportion of invariant sites, *p*_*invariant*_ to model functionally essential amino acids. Mutation follows the LG model of amino acid substitution ((Le and Gascuel 2008); Supplementary Figure 9), with transition probabilities defined by the matrix **L**, where *L*_*a,a*′_ is the probability that amino acid *a* transitions to amino acid *a*′, with *a* ≠ *a*^′^. Mutation probabilities vary across sites in the protein (defined by the vector **m**) calculated as the median values of four quartiles of 10 000 samples drawn from a gamma distribution after (Yang 1994) such that 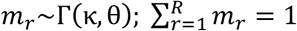 (Supplementary Figure 10). Within each generation, proteins crossing the stability threshold (*T* > Ω) and those randomly selected according to *p*_*death*_ are killed and randomly replaced *in populous* by a stable protein generating independent lineages, preventing star phylogenies.

Evolution is simulated with population isolation to mimic bifurcations. The protein population bifurcates every *g*_*B*_ generations into independent subpopulations (Φ_*branches*_) which undergo sequence replacement *in populus* (Supplementary Figure 11). The bifurcation interval is defined as 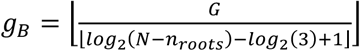 such that there are always 3, 4 or 5 proteins left in each branch at the end of the simulation. If every protein in a subpopulation is below the stability threshold, the simulation reverts to the prior generation to re-mutate the sequences and try to avoid complete branch extinction.

PESST continually tracks changes in stability at the amino acid (Δ_*r,a*_), protein (*T*) and population 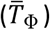 levels. From these recorded stabilities, animated figures are produced in addition to statistical analyses and FASTA files for ancestral reconstruction. The model is summarised in Algorithm 1 with additional symbols in Supplementary Table 1. The model was implemented using Python v3.8 (Van Rossum and Drake 2009) and the code is open source and freely available for download^3^.

#### Algorithm 1

PESST evolutionary algorithm pseudocode.

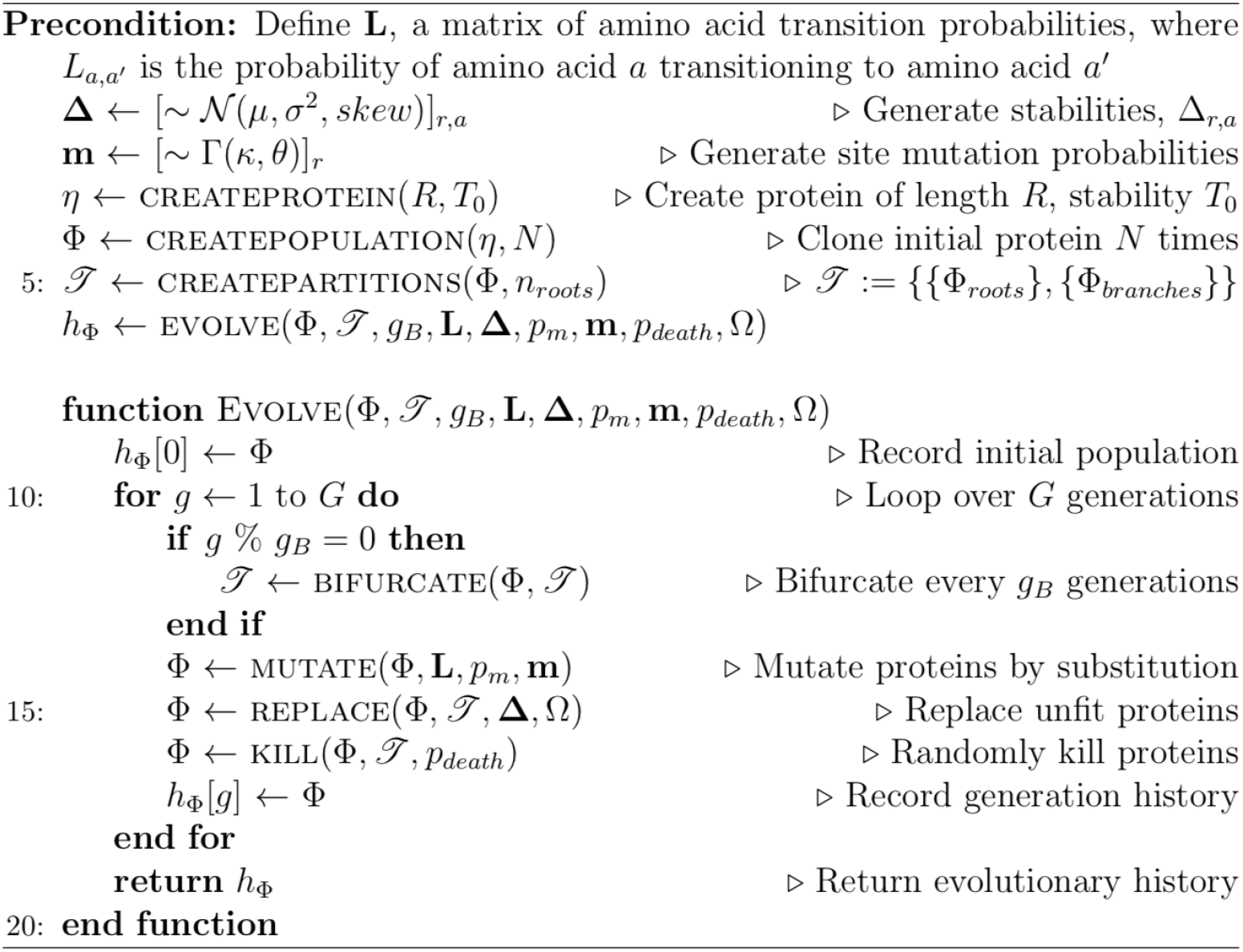

## Statistical tests

Equality comparisons of Δ_*r,a*_ distributions in the evolving dataset and the global stability matrix **Δ** were performed with the Kolmogorov-Smirnov test, computed by PESST. Equality meta-analyses were manually computed with Fisher’s combined probability analysis: 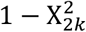 where 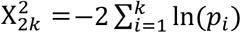 with 2*k* degrees of freedom, where *k* is the number of *p*-values from the 2-sided Kolmogorov-Smirnov tests in the meta-analysis; (Winkler et al. 2016). Analyses of the difference between the populations of ancestral and extant stabilities in each simulation were performed with both the Mann-Whitney *U* test and Welch’s *t*-test in Graphpad PRISM v9.2.0, as we cannot assume the normal distributions in every case. Analyses of correlations between node age and stability were computed for all replicate simulations in a given condition with Spearman’s method in Graphpad PRISM v9.2.0. Trees used for the analyses were cladograms output from CodeML, where age was defined as the number of nodes preceding a node of interest until the root is reached. Stabilities were the averaged normalized stability of ancestors in the longest subtrees containing at least three representative nodes of a given maximum subtree length. Equality comparisons between the stabilities of ancestors or consensus sequences of separate simulation conditions were performed with the Mann-Whitney *U* test in Graphpad PRISM v9.2.0.

### Ancestral Sequence Reconstruction

FASTA files were imported into the Geneious Prime sequence analysis suite (ver. 2022.1.1 (Kearse et al. 2012)). Phylogenies were produced with PhyML (Guindon et al. 2010) under standard settings with the LG+I+G model of amino acid substitution with estimated rates (Le and Gascuel 2008). SH-like branch supports were computed for phylogenies. Resulting phylogenies were manually rooted on the root sub-population defined by PESST. Marginal reconstruction of ancestors within the dataset was performed with CodeML of the PAML software suite (Yang 2007). Reconstructions of these data were performed under standard settings implementing the LG model of amino acid substitution with an estimation of gamma and of invariant sites (Le and Gascuel 2008). Both reconstructed sequences and PAML generated cladograms were extracted from PAML outputs. Reconstructed sequences’ stability values were calculated with `PESST.extras.fasta.process_fasta`, which cross references a user defined stability matrix with a FASTA formatted list of sequences of equal length. Output stabilities were analysed manually.

### Consensus sequences

Consensus sequences of FASTA formatted alignments were generated with `PESST.extras.fasta.write_concensus`. This is a low powered consensus sequence builder that generates up to three consensus sequences from a user input alignment by outputting the most common amino acid at every site. For alignments with ambiguous sites (multiple amino acids are equally common), the algorithm outputs up to three sequences built from the second and if present, third equally likely amino acids. Consensus sequence stability was calculated with `PESST.extras.fasta.process_fasta` as before. Output stabilities were analysed manually.

## Acknowledgements

This work was supported by a Biotechnology and Biological Sciences Research Council studentship (grant number BB/J014400/1) to AT; and a Wellcome Trust Institutional Strategic Support Award (grant number 204909/Z/16/Z) to BDE.

## Footnotes

1 PESST is written in Python and freely available under version control at https://github.com/bdevans/PESST.

2 *p*-values from Kolmogorov-Smirnov tests ranged from 3.4 × 10^−155^ to 5.1 × 10^−186^.

3 https://github.com/bdevans/PESST

