## Supplementary Figures and Tables for "Survivor bias drives overestimation of stability in reconstructed ancestral proteins"

**Supplementary Figure 1**

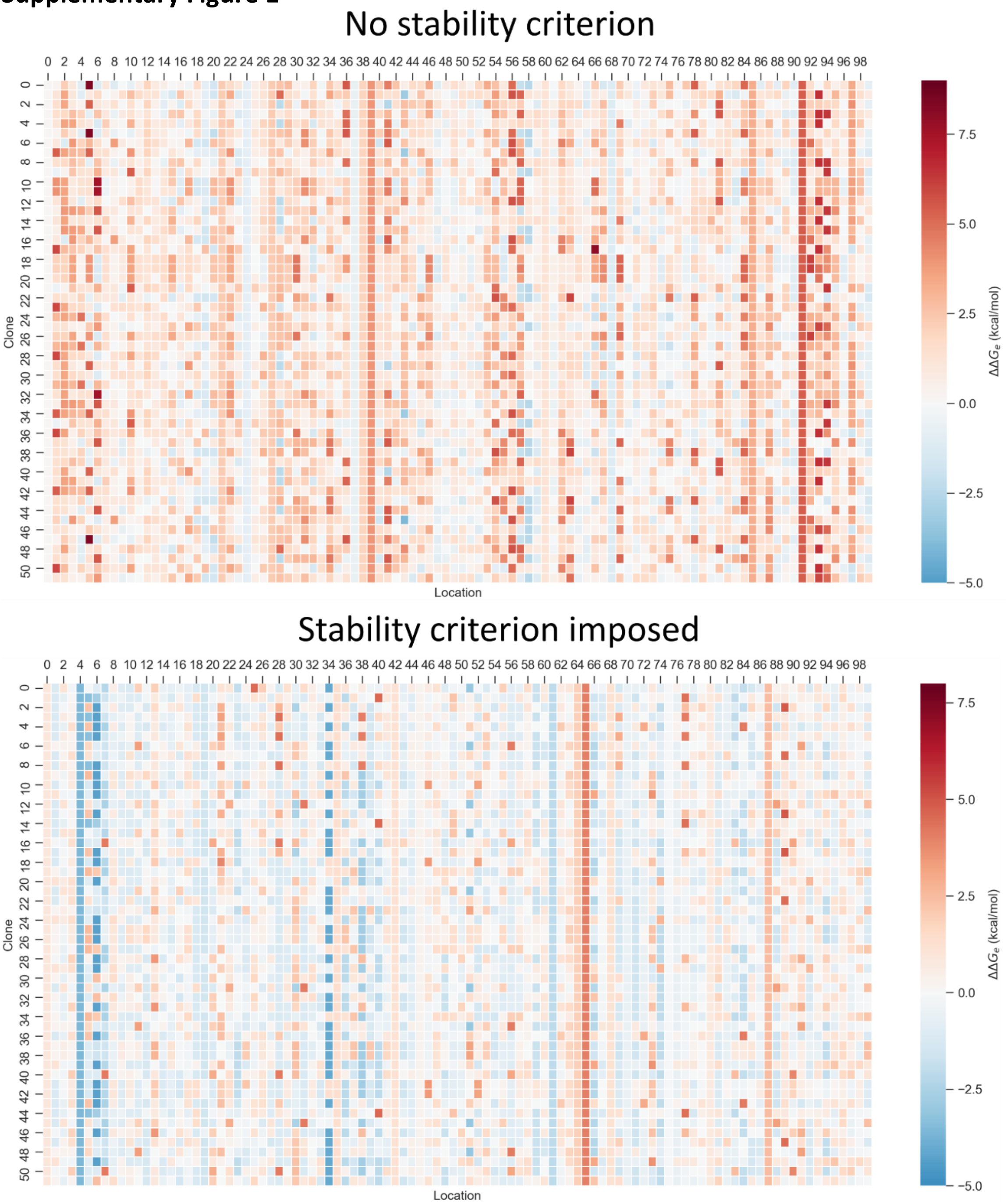


**Supplementary Figure 1 – Imposing a stability criterion results in a shift to more stabilising locus $\Delta G_0$ values.**

Images show heat maps of locus $\Delta G_0$ values from simulations that do not (upper panel; same simulation as Figure 1E) or do (lower panel; same simulation as Figure 2) impose a fitness criterion. Heat maps were generated at generation 5 000. Populations that have evolved under a selective pressure for fitness have considerably more stabilizing residues than populations released from this selective pressure. Stabilizing residues are therefore overrepresented in populations evolving with the selective pressure of a stability criterion.

**Supplementary Figure 2**

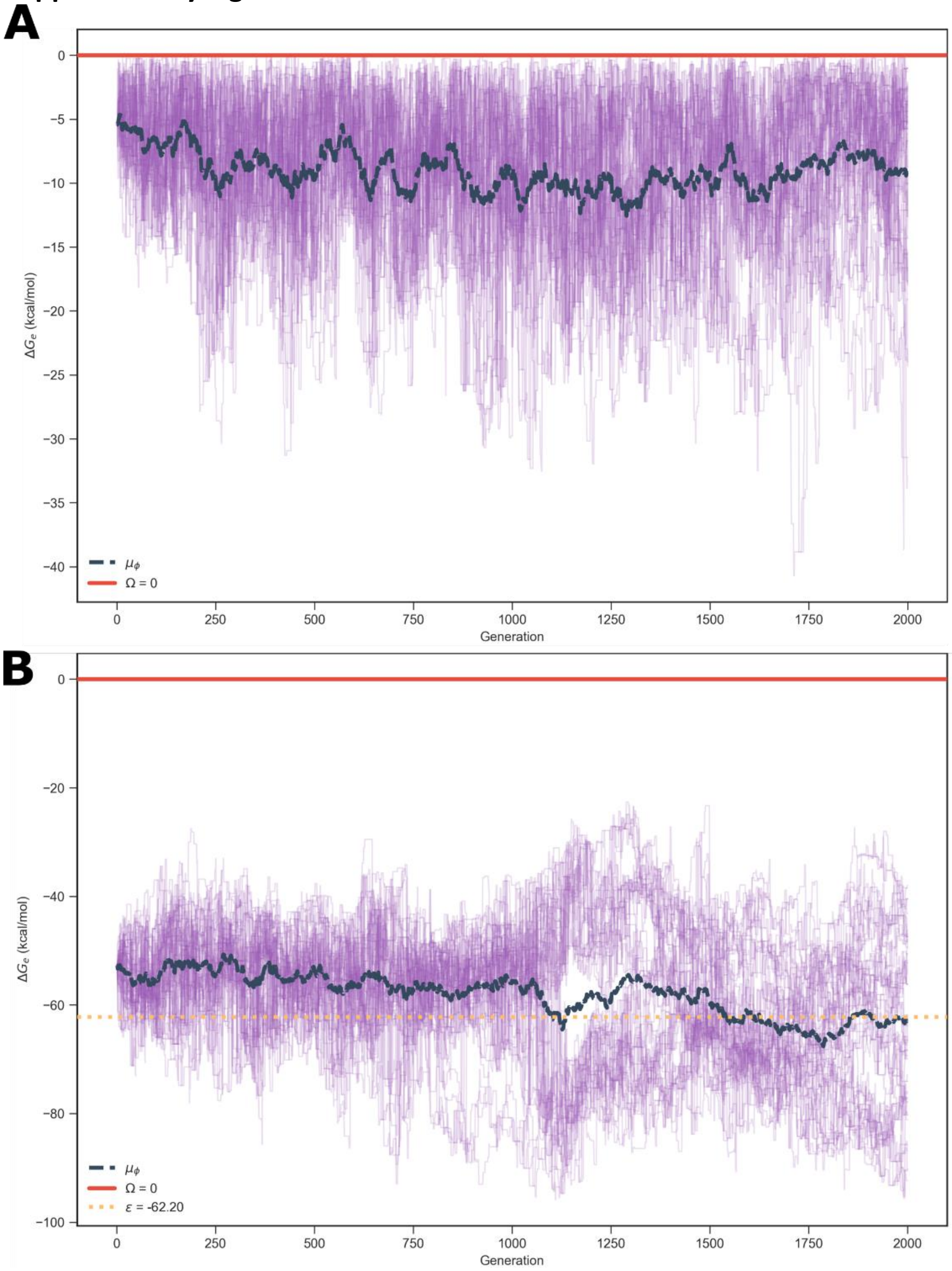


**Supplementary Figure 2 – Simulations with a synthetic global stability matrix without a negative average explore a wide stability space due to release of selective pressure.**

Representative stability traces for PESST simulations of 2 000 generations where the global stability matrix is not destabilising. Simulated data used the natural stability distribution with the mean values artificially set to zero (**A**) or -0.5x the experimentally determined values (**B**). With reduced or no selective pressure, the population broadly samples stability space. Individual lines represent the stability of one of 52 clones in the dataset, which are each tracked independently and simultaneously by PESST. The solid red line represents the stability threshold Ω. The dashed black bold line represents the average stability of the population. Data shown are for the first replicate for experiments **4c** and **4d**.

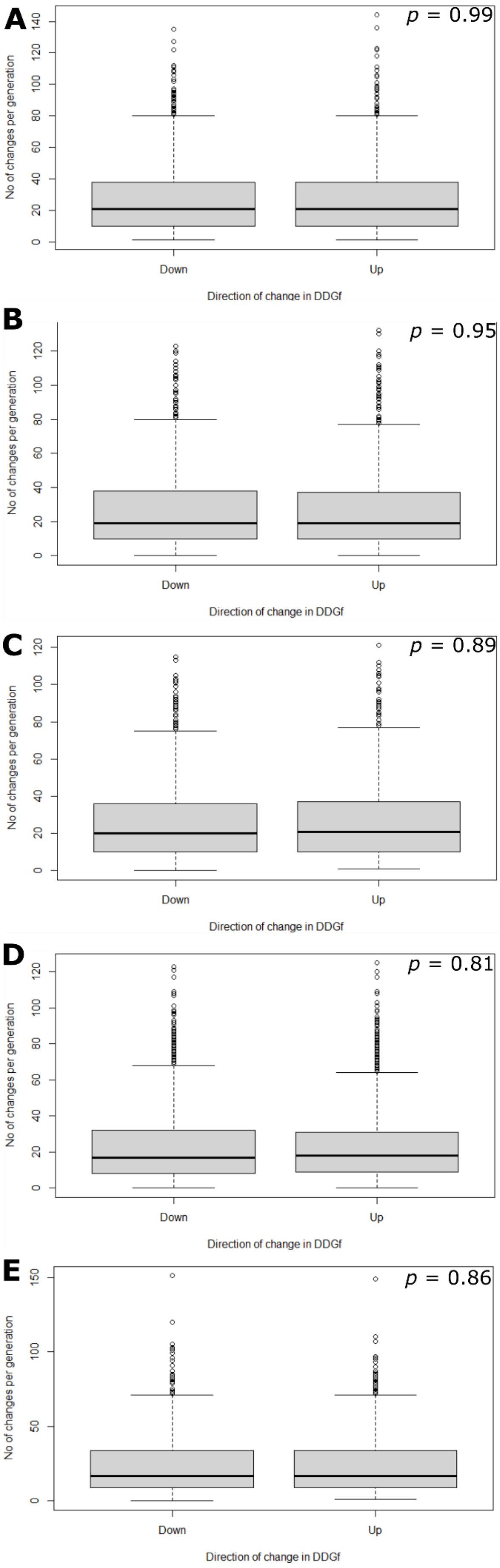


**Supplementary Figure 3**

**Supplementary Figure 3 – Mutations that increase and decrease stability are sampled equally.**
Changes in amino acid sequence were followed across 1 750 generations for simulations with a stability criterion enforced (data from experiment **4a**). The first 250 generations were not tracked as these are still coming to equilibrium. For each generation the number of mutations that increase and decrease stability was tracked. The total number of positive and negative changes per generation are plotted for each replicate. For each replicate, a *t*-test was performed (shown on figures). There was no significant difference between positive and negative changes in any replicate. The analysis was performed using RStudio v. 2022.02.2, and the code used is supplied in the supplementary files (mutation_check.R).

**Supplementary Figure 4**

**A** Natural stability distribution

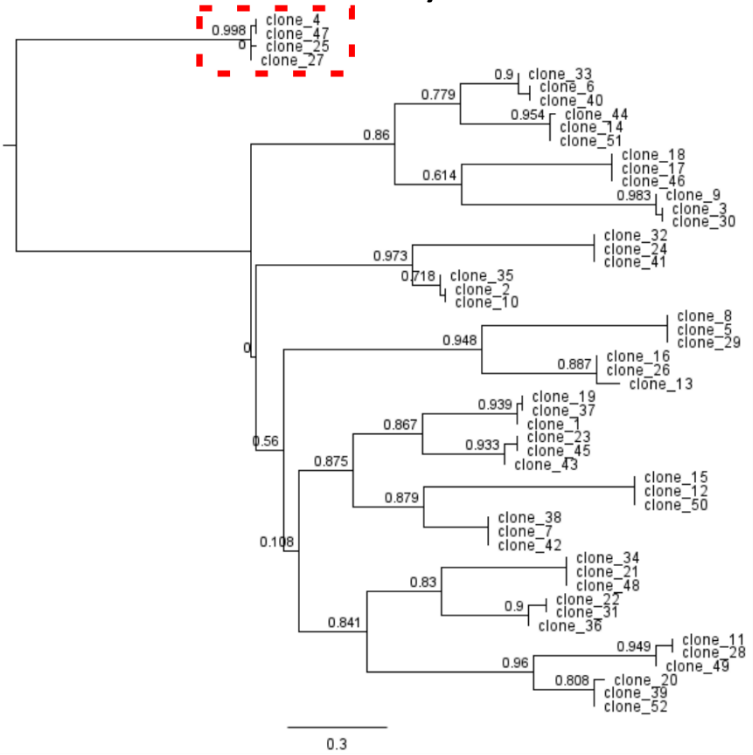


Stability matrix mean of 0.5x natural mean

**B**

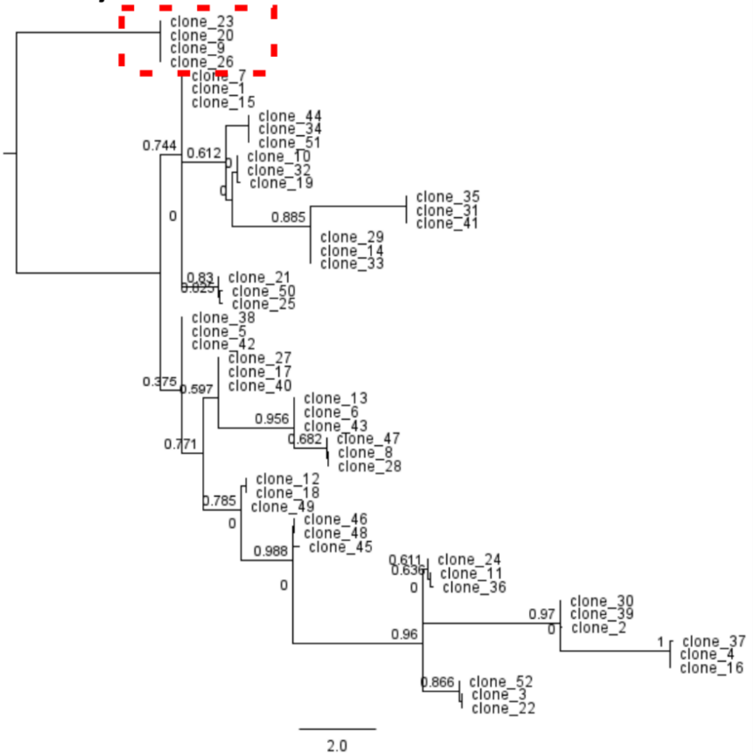

**C** Stability matrix mean zero

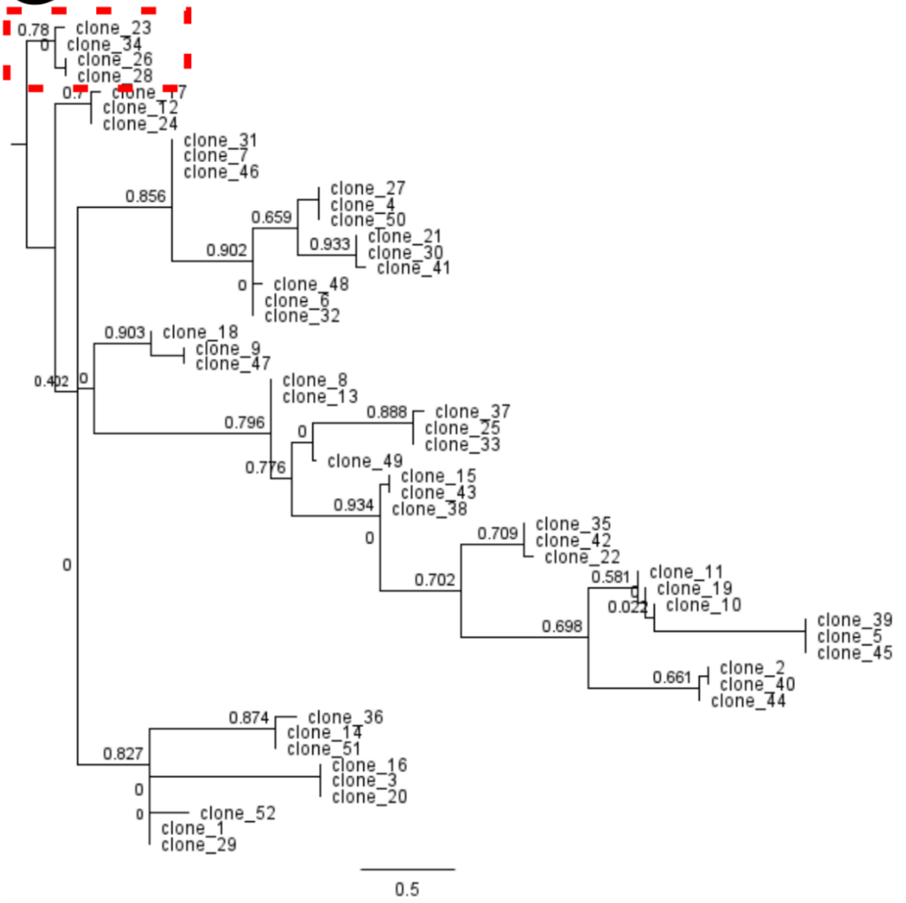


**D** Stability matrix mean of -0.5x natural mean

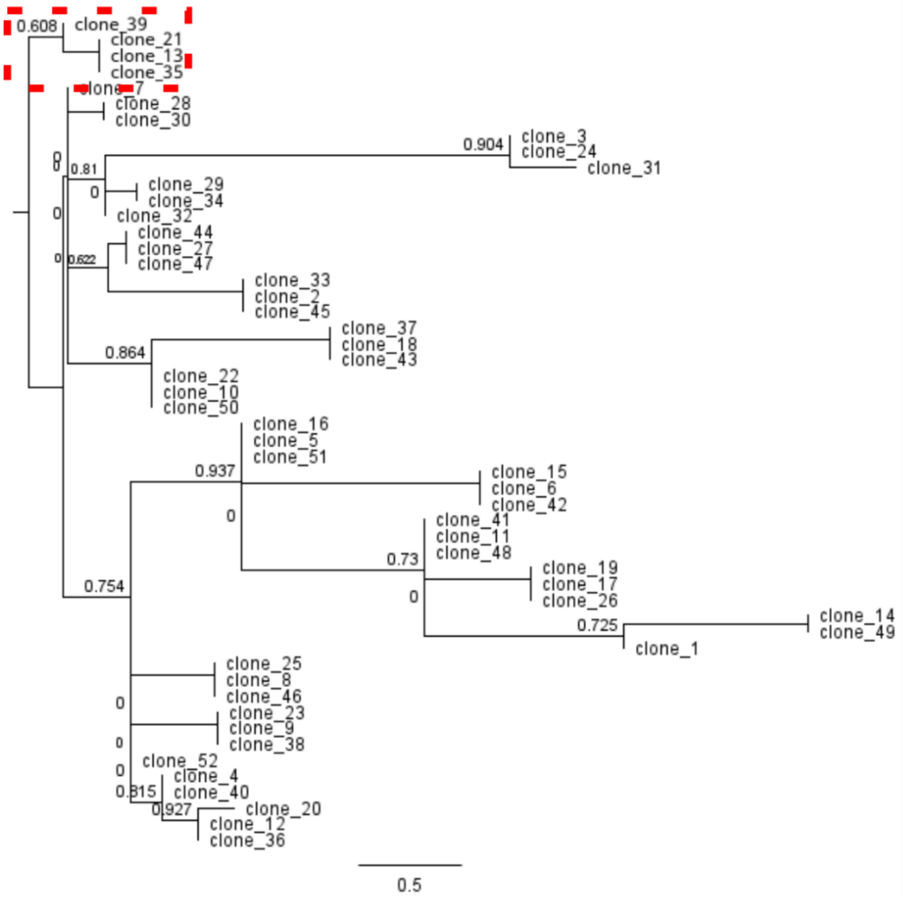

**Supplementary Figure 4 – Representative phylogenies generated from PESST simulations used for Figures 3 and 4.**

Alignments were generated from PESST simulations of 2 000 generations where the global stability matrix is not destabilising. Simulated data used the natural stability distribution with the natural mean (**A**), or mean values artificially set 0.5x (**B**), 0x (**C**) or -0.5x (**D**) the experimentally determined values. Phylogenies of simulated evolution were generated with PhyML (Guindon *et al.,* 2010). Data presented is representative of single simulations under the given conditions (Run 1 used for **A, B, C**; Run 2 used for **D** as Run 1 showed a single unusual group that made visualisation of the remaining data challenging). Phylogenies were modified in Geneious version 2022.1 (Biomatters, available from https://www.geneious.com). Node values represent SH-like support values calculated in PhyML. Scale represents mutations per site.

**Supplementary Figure 5**

**A**

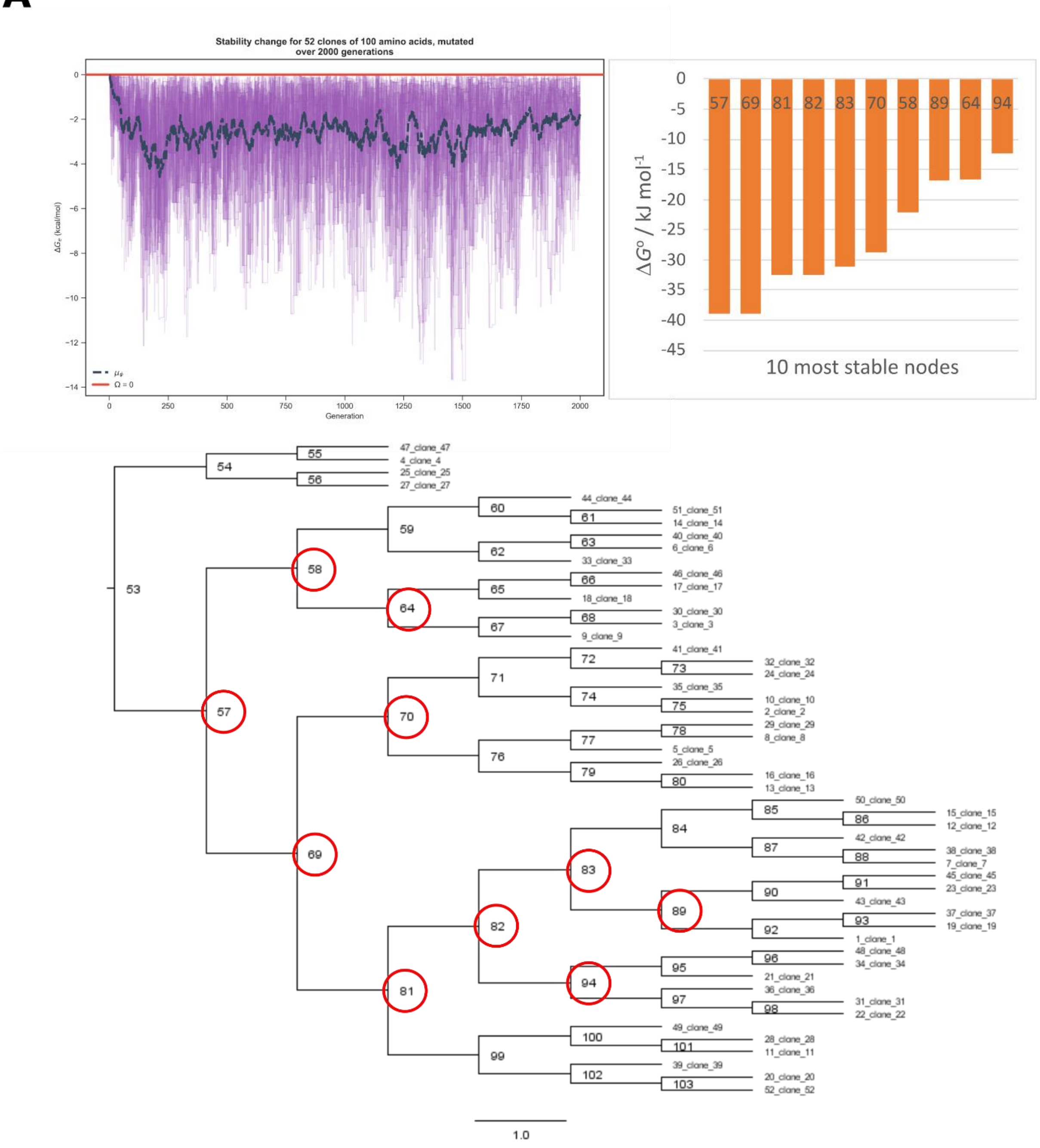

**B**

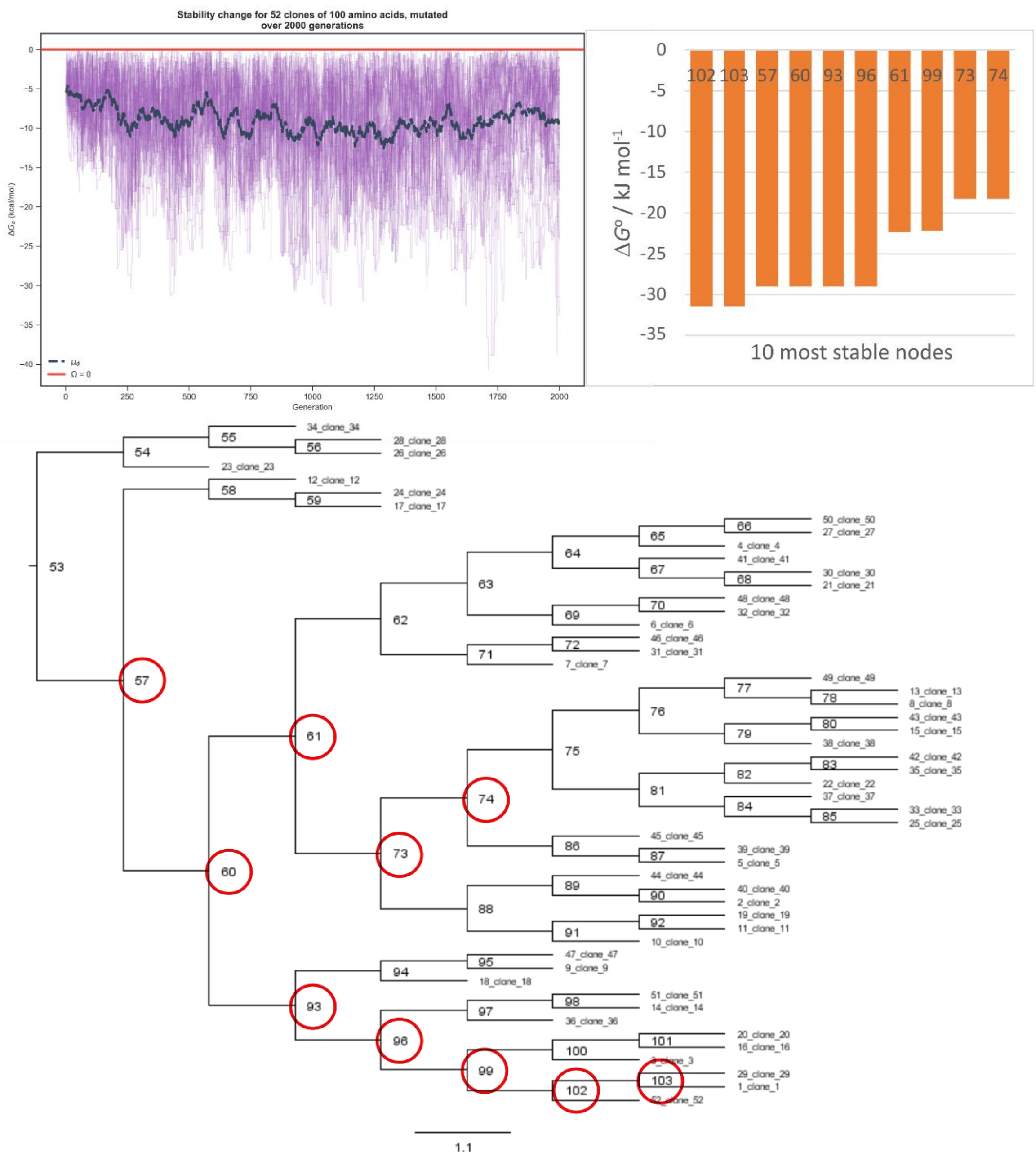


### Supplementary Figure 5 – Representative simulations showing that reconstructed ancestors explore a higher stability space than that of their evolutionary history when a bidirectional selective pressure is present.

Data representative of ancestral sequences generated from PESST simulations of 2 000 generations with a fitness criterion imposed. Simulated data used the natural stability distribution with the natural mean (**A**), or mean values artificially set to 0x (**B**). **Top left**: Stability trace from the representative simulation. **Top right**: Stability of the ten most stable ancestral nodes derived from a given simulation calculated with CodeML in PAML (Yang 2007). **Bottom**: Cladogram output by PAML visualised in FigTree (Rambaut 2018). Node labels represent the ancestor sequence identifier produced by PAML, and the top ten nodes are highlighted with red circles. Ancestral reconstruction of simulations evolving under a bidirectional selective pressure (**A**) leads to proteins with stabilities that are higher than any stability sampled by the evolving population. This effect does not occur when the bidirectional selective pressure is released (**B**).

### Supplementary Figure 6

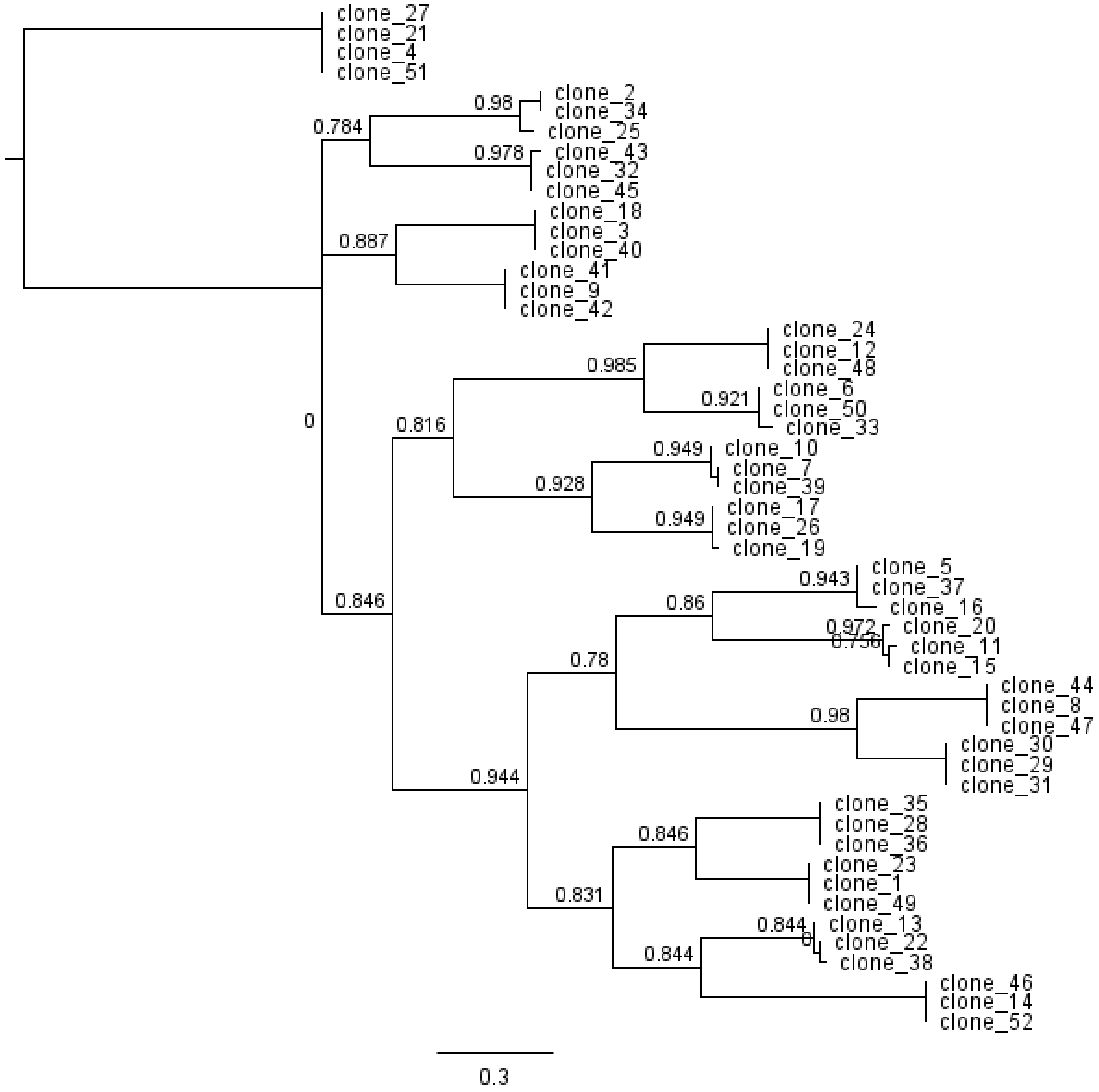


**Supplementary Figure 6 – Phylogeny for simulation that evolves from high stability to a threshold.**

PESST simulations were initiated at high stability (random choice of the three most stabilising amino acids in each evolving position). The alignments generated were exported into Geneious ver. 2022.1 (Biomatters, available from https://www.geneious.com) (Kearse et al. 2012) . Phylogeny of simulated evolution was generated with PhyML (Guindon et al. 2010). Data presented is representative of a single simulation under the given conditions. Data for all simulations is available in supplementary files. The phylogeny was visualised in Geneious version 2022.1. Node values represent SH-like support values calculated in PhyML. Scale represents mutations per site.

**Supplementary Figure 7**

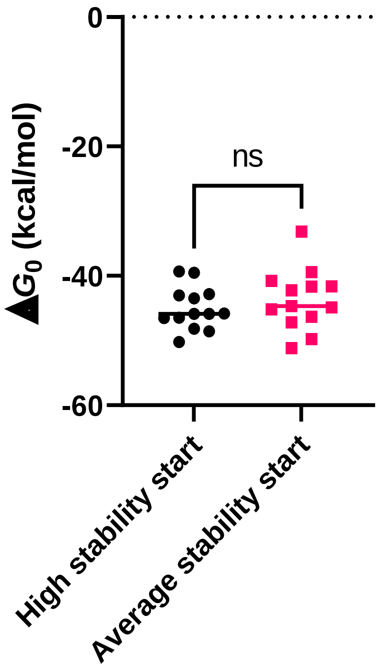


**Supplementary Figure 7 – Consensus sequences from simulations initiated at high stability show similar stability to those initiated at average stability.**

Grouped min-max scatter of the stabilities of consensus values derived from the five simulations of both scenarios using used the natural stability distribution with the natural mean. The figure compares sequences evolving from a starting point of high stability (black circles) or average stability (pink squares). The Mann Whitney *U* test was used to calculate whether the populations were significantly dissimilar ($p$ = 0.39).

### Supplementary Figure 8

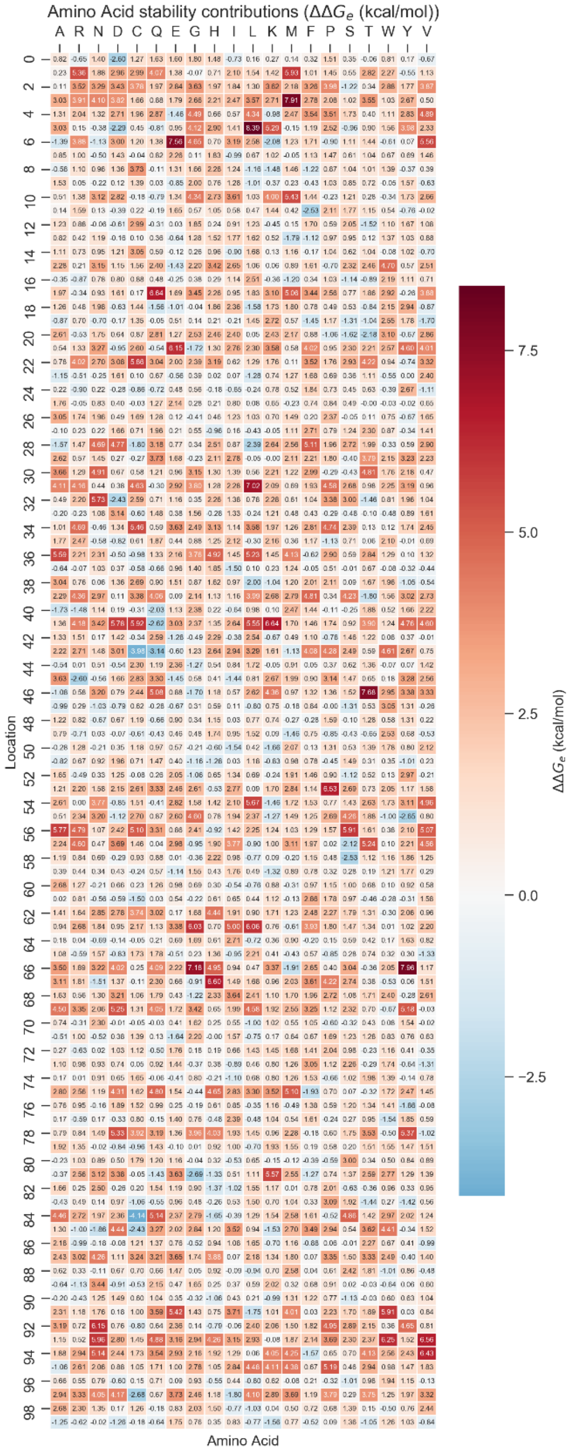


### Supplementary figure 8 – Representative $\Delta$ matrix generated in PESST simulations.

Heat map of a representative $\Delta$ matrix generated from a Gaussian distribution at the initiation of a PESST simulation. Each possible amino acid at each position of the protein is assigned a $\Delta_{r,a}$ value. The matrix remains consistent for the entire simulation and can be used to back-calculate the stabilities of protein sequences derived from the representative simulation. Given matrix is derived from the distribution from (Tokuriki et al. 2007).

**Supplementary Figure 9**

**A**

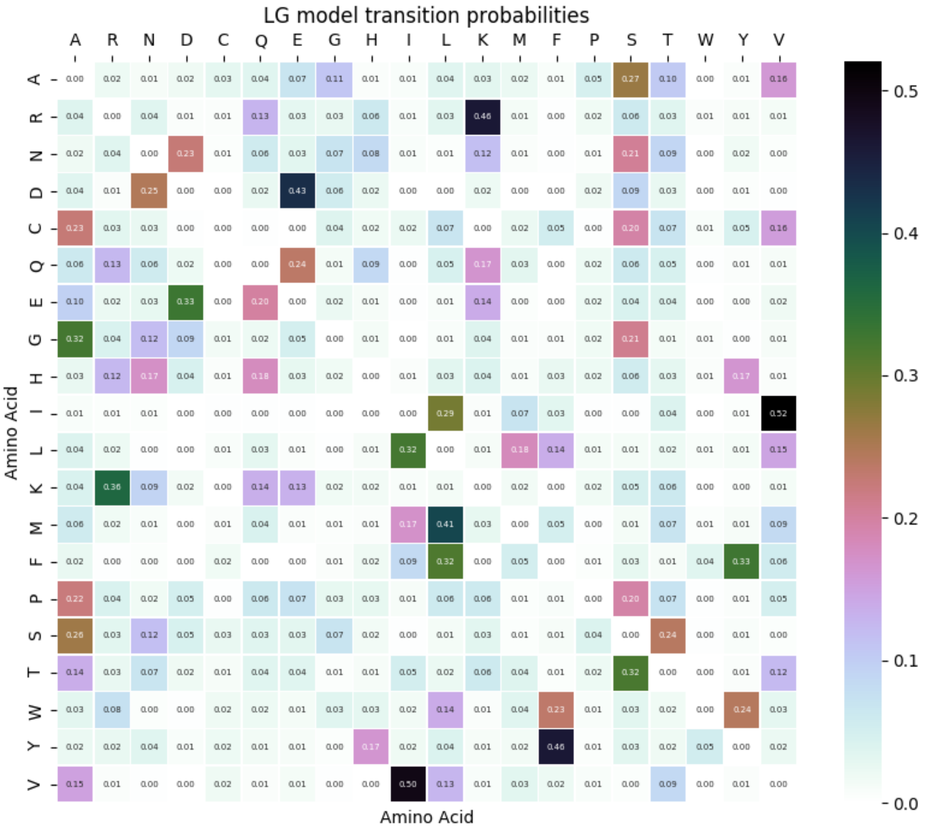


**B**

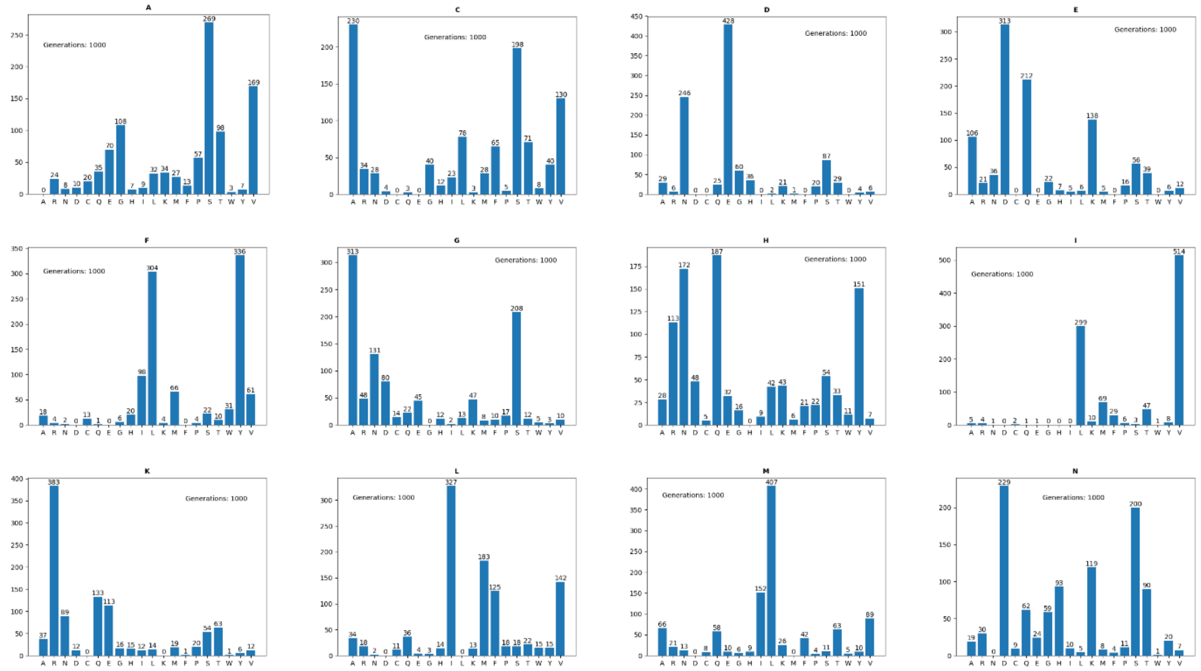

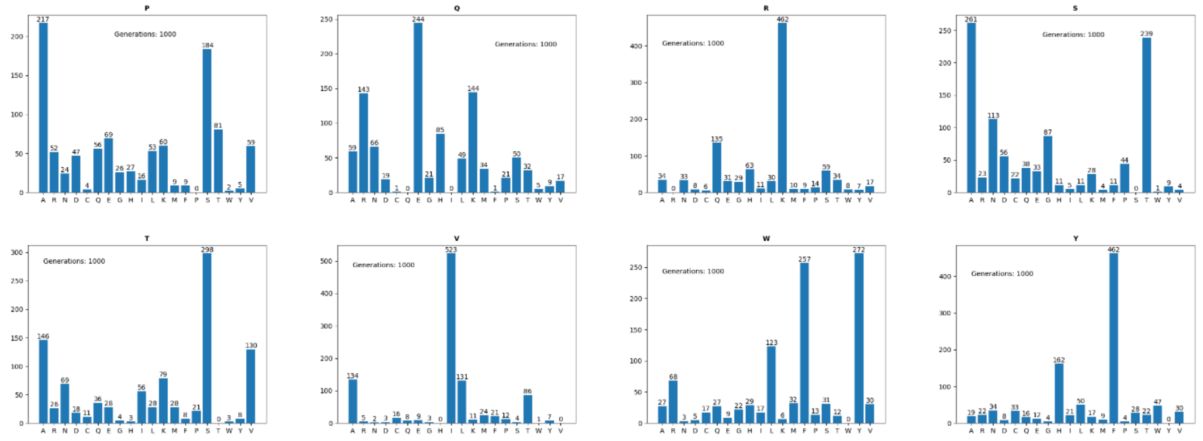


**Supplementary Figure 9 – Transition rates at each site derived from a modified LG model implemented into PESST.**

The LG model is a matrix of substitution likelihoods used to compute the likelihood and nature of mutations along a phylogenetic branch. Diagonals in the matrix describe the rate at which an amino acid does not change. By modelling evolution to a uniform clock and dictating that a set number of amino acid conversions must happen at each generation, the diagonal rates in the original LG model become moot. **A**: The LG matrix with diagonal values set to 0 and the remaining values normalised to 1 providing a means to calculate the substitution likelihoods when an amino acid change is forced. **B:** The distribution of substitutions that occur according to the model from 1 000 generations of mutation at each possible amino acid state, suggesting the LG model is implemented as expected.

**Supplementary Figure 10**

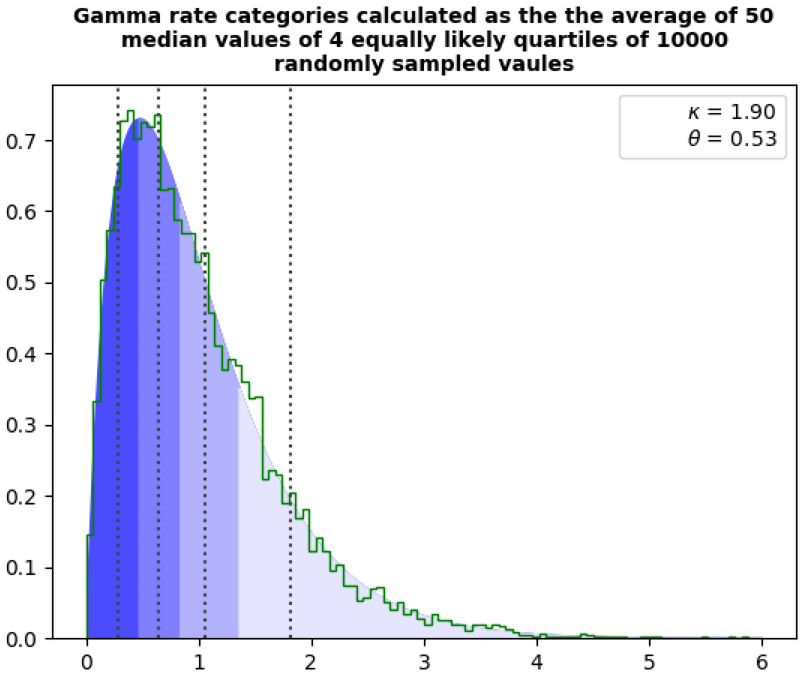


**Supplementary Figure 10 – Four independent rate categories defined by the median values of four quartiles from 10 000 samples of the gamma distribution.**

Gamma distribution sampling implemented into PESST. Across a protein, mutation rates are found to fit a gamma distribution, which can be simplified for computational purposes without cost to accuracy by assigning residues a mutation rate from one of four independent gamma rate categories. Rate categories are calculated as the median value of quartiles from 10 000 random samples of a gamma distribution.

### Supplementary Figure 11

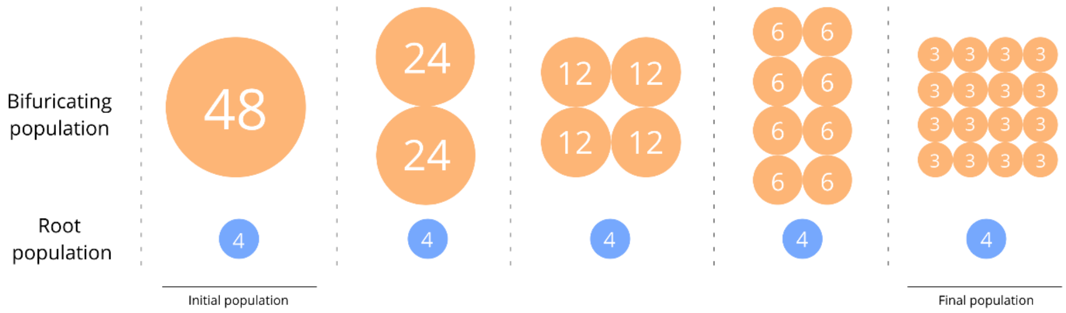


**Supplementary Figure 11 – implementation scheme of bifurcations in PESST**.

Scheme representing how bifurcations are implemented in PESST. Bifurcating the population "geographically separates" the data into subpopulations, meaning that as the protein population evolves, each sub-population can only replicate *in populous*. This is intended to be a rough mimic for natural bifurcating populations. At even generation intervals, the population will bifurcate completely once. The number of bifurcations is dependent on the population size. Bifurcations occur until all final sub-population sizes are 3, 4 or 5. Additionally a root sub-population is isolated from the data at generation 0. Implementation of bifurcation into PESST allows for increased phylogenetic signal within simulations, and the generation of clear well supported phylogenies.

**Supplementary Table 1**

| Parameter | Symbol | Notes |
|---|---|---|
| Population of proteins | $\Phi$ | $\{\eta_1, \eta_2, \cdots, \eta_N\}$ |
| Protein sequence | $\eta$ | $[a_1, a_2, \cdots, a_R]$ |
| Matrix of transition probabilities | $\mathbf{L}$ | $L_{a,a'} := p(a \rightarrow a'); a \neq a'$ |
| Matrix of amino acid stabilities | $\boldsymbol{\Delta}$ | |
| Amino acid stability contribution | $\Delta_{r,a}$ | |
| Protein stability | $T$ | $:= \sum_{r=1}^{R} \Delta_{r,a_r}$ |
| Mean protein stability | $\overline{T}_\Phi$ | $:= \frac{1}{N} \sum_{n=1}^{N} T_n$ |
| Vector of mutation probabilities | $\mathbf{m}$ | $m_r \sim \Gamma(\kappa, \theta); \sum_{r=1}^{R} m_r = 1$ |
| Evolutionary history | $h_\Phi$ | $[\Phi_1, \Phi_2, \cdots, \Phi_G]$ |
| Expected stability convergence | $\epsilon$ | $\approx \mu \cdot R$ |
| Population of roots | $\Phi_{roots}$ | $\{\eta_1, \eta_2, \cdots, \eta_{n_{roots}}\}$ |
| Population of branches | $\Phi_{branches}$ | $\{\{\eta_i\}_1, \{\eta_j\}_2, \cdots, \{\eta_k\}_{n_{branches}}\}$ |

**Supplementary Table 1 – Additional symbols used in PESST.** $\Delta_{r,a}$ is referred to as "locus $\Delta\Delta G$" in the main text.

### Supplementary Table 2

| Starting parameter | Simulation | Seed |
|---|---|---|
| High stability | 1 | 88340051 |
| | 2 | 2029880479 |
| | 3 | 1396291103 |
| | 4 | 924280734 |
| | 5 | 2602377145 |
| Average stability | 1 | 3582916783 |
| | 2 | 31087996 |
| | 3 | 3568441 |
| | 4 | 1461465320 |
| | 5 | 3582063969 |
| Low stability | 1 | 571460824 |
| | 2 | 2832642253 |
| | 3 | 3605296218 |
| | 4 | 2953479558 |
| | 5 | 852485793 |

### Supplementary Table 2 – Seeds for the simulations used for Figure 1.

PESST implements a pseudo random number generator to define various simulation variables, so each simulation represents a unique evolutionary scenario. To ensure that PESST simulations are replicable, the pseudo random number generator can be seeded. The seeds used for the simulations that make up the results in Figure 1 are presented.

**Supplementary Table 3**

| Starting parameter | Simulation | Seed |
|---|---|---|
| High stability, stability criterion imposed | 1 | 3132065010 |
| | 2 | 2569891490 |
| | 3 | 3669476248 |
| | 4 | 510649927 |
| | 5 | 4172610431 |

**Supplementary Table 3 – Seeds for the simulations used for Figure 2.**

### Supplementary Table 4

| Starting parameter | Simulation | Seed |
|---|---|---|
| Average stability<br>Stability criterion<br>Natural Global stability matrix | 1 | 453390725 |
| | 2 | 380194671 |
| | 3 | 2388428319 |
| | 4 | 199479308 |
| | 5 | 2589354605 |
| Average stability<br>Stability criterion<br>Global stability matrix with mean set to 0.5x natural mean | 1 | 1851761159 |
| | 2 | 2781027769 |
| | 3 | 2811992013 |
| | 4 | 2382803527 |
| | 5 | 3420674386 |
| Average stability<br>Stability criterion<br>Global stability matrix with mean set to zero | 1 | 3026563474 |
| | 2 | 2310734369 |
| | 3 | 4288522741 |
| | 4 | 79415063 |
| | 5 | 3333796942 |
| Average stability<br>Stability criterion<br>Global stability matrix with mean set to -0.5x natural mean | 1 | 1693768349 |
| | 2 | 3300881123 |
| | 3 | 116399059 |
| | 4 | 2412070430 |
| | 5 | 630218252 |
| High stability<br>Stability criterion<br>Natural Global stability matrix | 1 | 2145210219 |
| | 2 | 670959984 |
| | 3 | 3074709133 |
| | 4 | 2099671525 |
| | 5 | 1577985694 |

**Supplementary Table 4 – Seeds for Figures 3 and 4.**

### Supplementary Table 5

| $\overline{\Delta_{r,a}}$ | -2 | | -1 | | 0 | | 1 | |
|---|---|---|---|---|---|---|---|---|
| Test | Mann-Whitney U (*U*, *p*) | Welch's t-test (*t*, *p*) | Mann-Whitney U (*U*, *p*) | Welch's t-test (*t*, *p*) | Mann-Whitney U (*p*) | Welch's t-test (*p*) | Mann-Whitney U (*p*) | Welch's t-test (*p*) |
| **Simulation 1** | 879, 0.0029 | 3.67, 0.0005 | 950, 0.0126 | 2.99, 0.0039 | 1145, 0.2329 | 0.98, 0.3285 | 1133, 0.2045 | 1.13, 0.2596 |
| **Simulation 2** | 888, 0.0036 | 3.82, 0.0004 | 962, 0.0158 | 3.82, 0.0003 | 1133, 0.2033 | 1.11, 0.2664 | 1200, 0.4065 | 0.61, 0.5454 |
| **Simulation 3** | 837, 0.0011 | 3.89, 0.0003 | 911, 0.0059 | 3.31, 0.0015 | 1155, 0.2612 | 0.96, 0.3419 | 1203, 0.4197 | 0.49, 0.4865 |
| **Simulation 4** | 789, 0.0003 | 4.16, 0.0001 | 855, 0.0017 | 3.38, 0.0013 | 1179, 0.3328 | 1.06, 0.2907 | 1138, 0.2153 | 1.68, 0.0973 |
| **Simulation 5** | 948, 0.0121 | 2.94, 0.0044 | 740, <0.0001 | 4.25, <0.0001 | 1056, 0.0751 | 2.18, 0.0319 | 1075, 0.0976 | 2.01, 0.0471 |

**Supplementary Table 5 – Simulation *p*-values in Figure 3.**

Table represents statistics and confidence values (*p*-values) obtained with the Mann-Whitney U test* or Welch's t-test when comparing the similarity of the distribution of comparing the stability values of proteins generation 2 000 to PAML calculated ancestors of the simulation. PESST simulations were initiated at $T^{\Omega+5;\Omega+25}$,where $p_m = 0.002$, $\Omega = 0$, and $\overline{\Delta_{r,a}} = -2, -1, 0$ and 1. Distributions of $T_m$ values that are not significantly different ($p > 0.05$) are coloured red.

* For all tests, $U_{Max}$ = 2 652. $U_{max}$ is defined as $n_1n_2$
