## Supplementary material for "Survivor bias drives overestimation of stability in reconstructed ancestral proteins": mutation_check.R

```
# Script to interrogate results from PESST
# Aim is to discover whether stabilising mutations are common or rare

# Load packages (if necessary)

# Set directories
setwd("C:/pesst/results/22-03-29-15-06/data")

Total <- data.frame(Direction = c(rep("Up",1750),rep("Down",1750)),
                    Number = c(0))
stab_01 <- matrix(rep(0,5100),nrow = 51, ncol = 100, byrow = TRUE)


for (gen in 1:1750){
  stab0 <- read.csv(paste0("stabilities_G",gen+249,".csv"))
  stab1 <- read.csv(paste0("stabilities_G",gen+250,".csv"))
  for (i in 1:51){
    up <- 0 ; down <- 0
    for (j in 1:100){
      stab_01[i,j] <- stab1[i,j] - stab0[i,j]
      if (stab_01[i,j] < 0) { down <- down + 1}
      if (stab_01[i,j] > 0) { up <- up + 1}
    }
    #if (up + down > 10) {up <- 0; down <- 0}
    Total$Number[gen+1750] <- Total$Number[gen+1750] + down
    Total$Number[gen] <- Total$Number[gen] + up
  }
}

boxplot(Total$Number~ Total$Direction, ylab="No of changes per generation", xlab="Direction of change in DDGf")
t.test(Total$Number~ Total$Direction, paired = FALSE)
```
