## Supplementary figures and images for "Survivor bias drives overestimation of stability in reconstructed ancestral proteins"

### Fig4a1.png

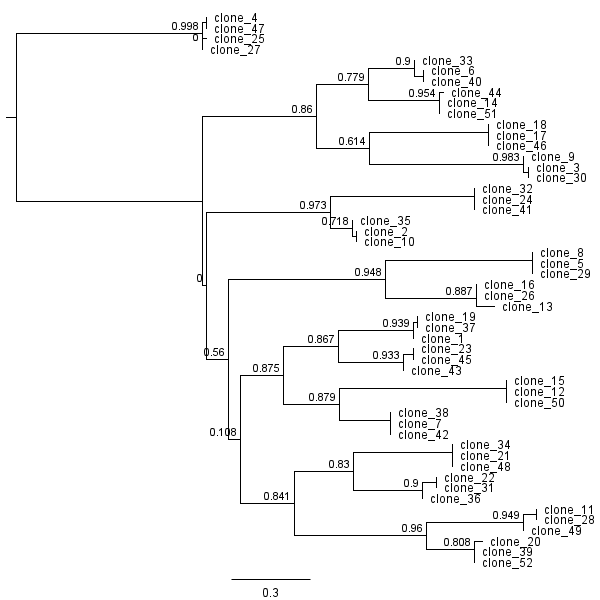

### Fig4a2.png

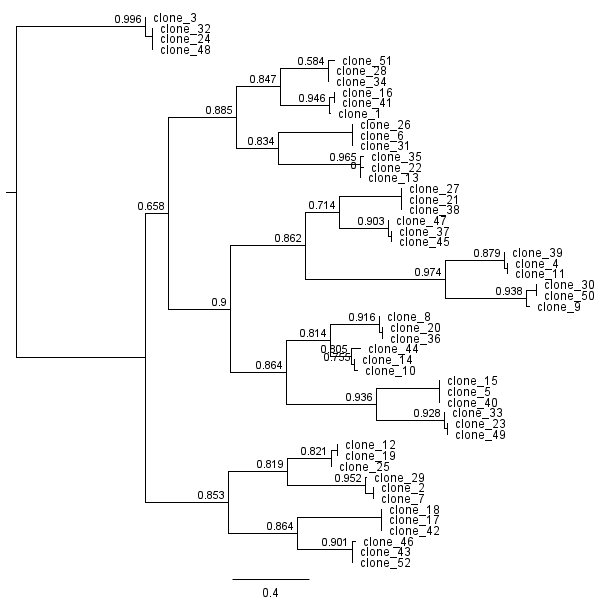

### Fig4a3.png

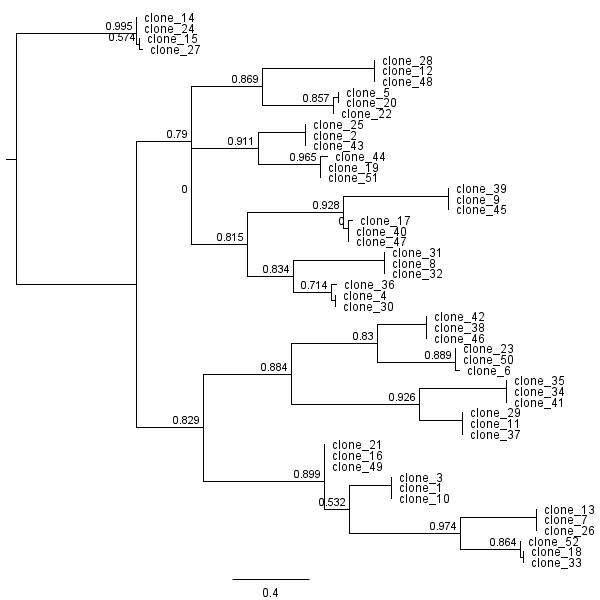

### Fig4a4.png

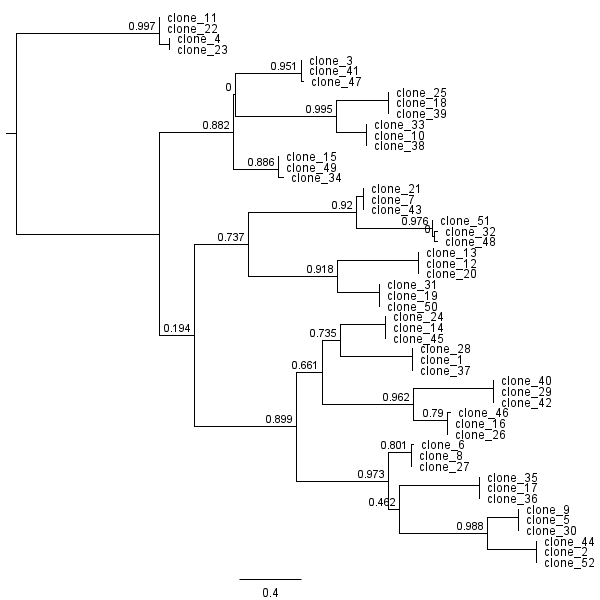

### Fig4a5.png

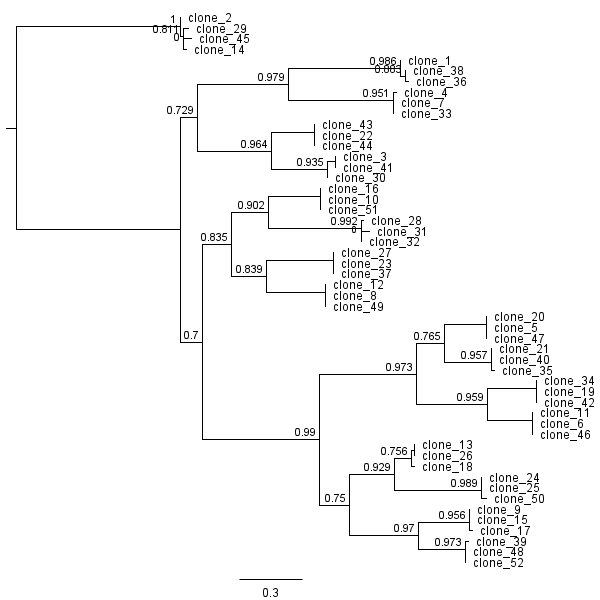

### Fig4b1.png

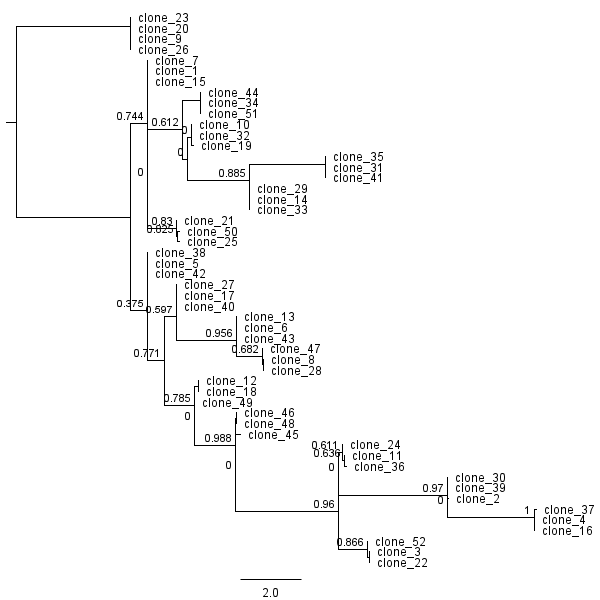

### Fig4b2.png

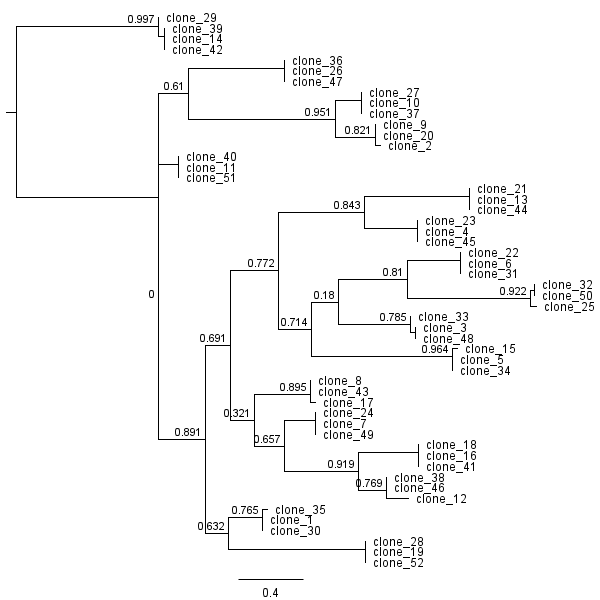

### Fig4b3.png

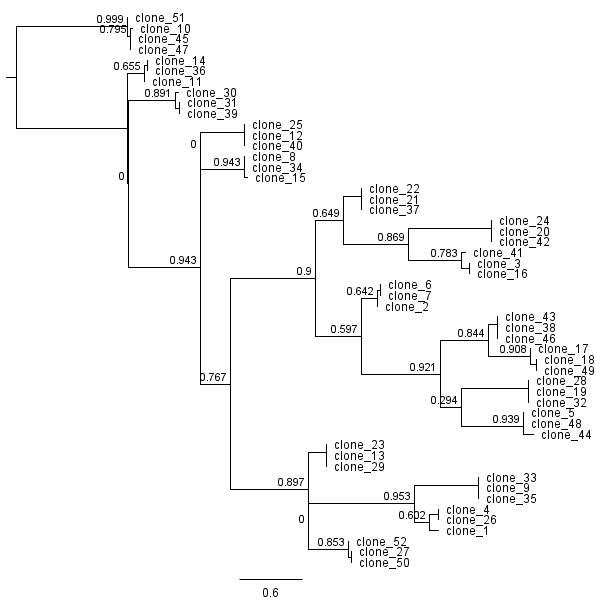

### Fig4b4.png

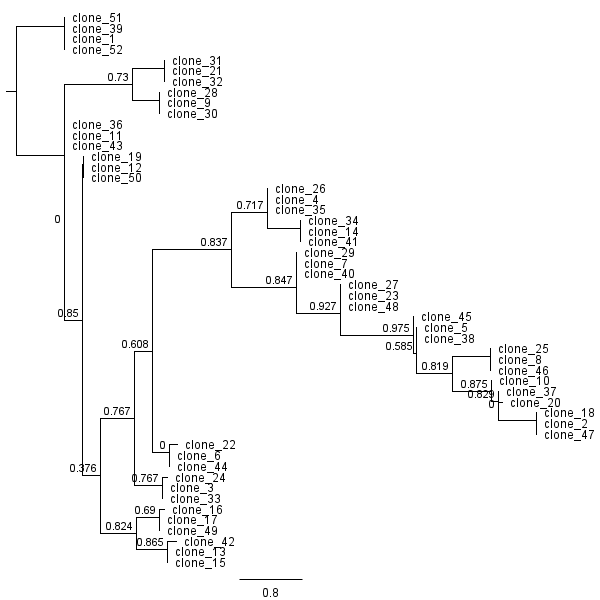

### Fig4b5.png

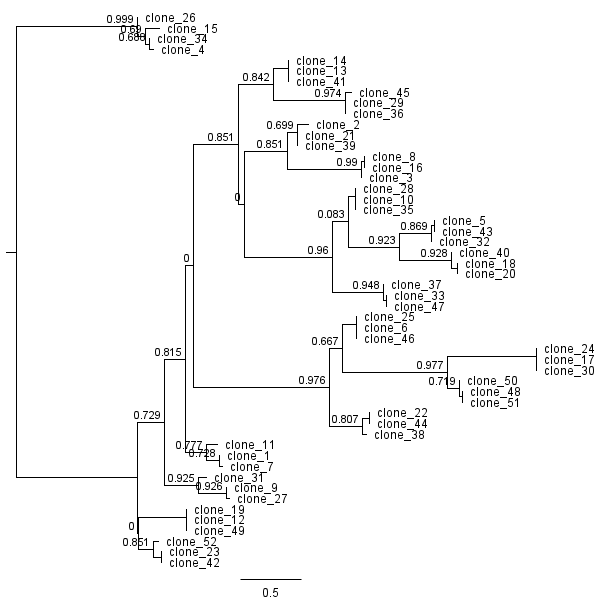

### Fig4c1.png

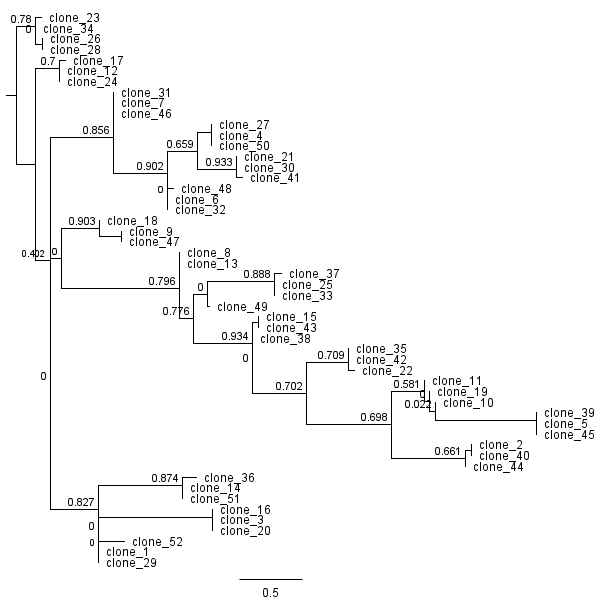

### Fig4c2.png

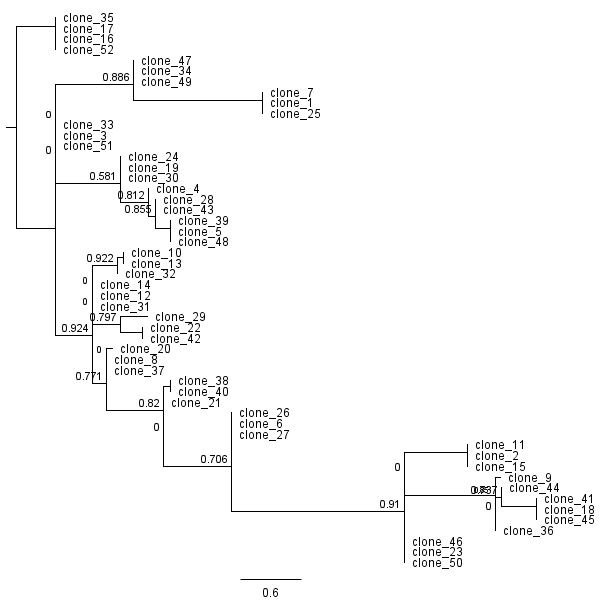

### Fig4c3.png

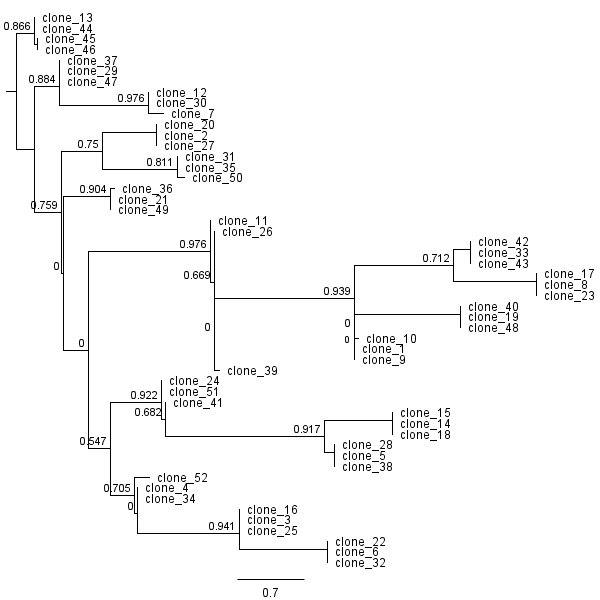

### Fig4c4.png

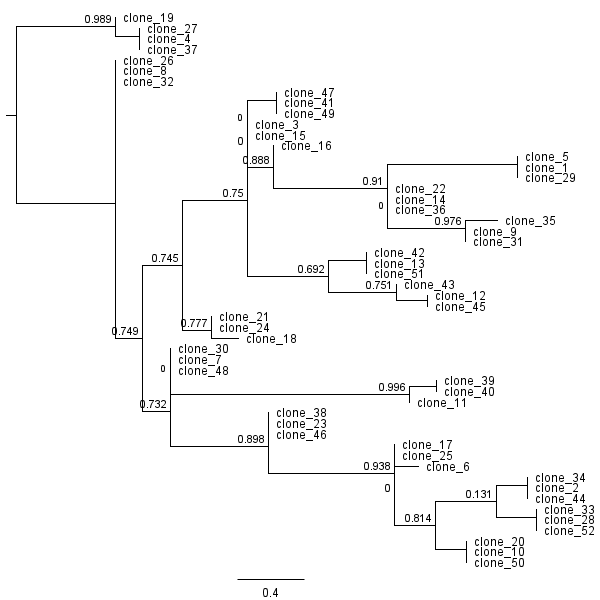

### Fig4c5.png

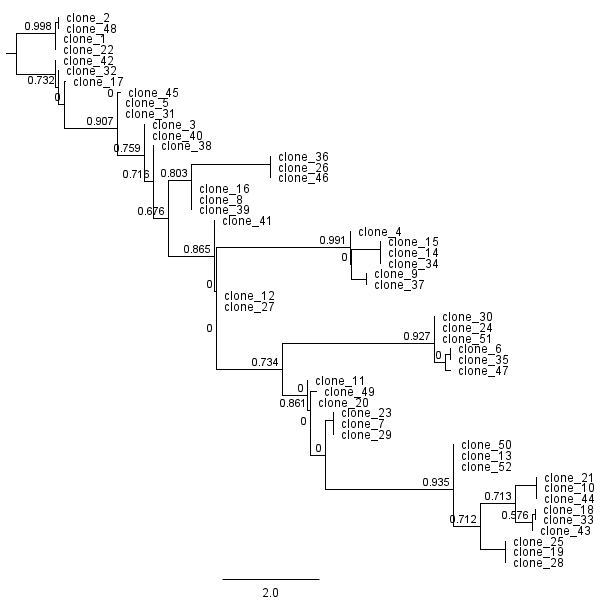

### Fig4d1.png

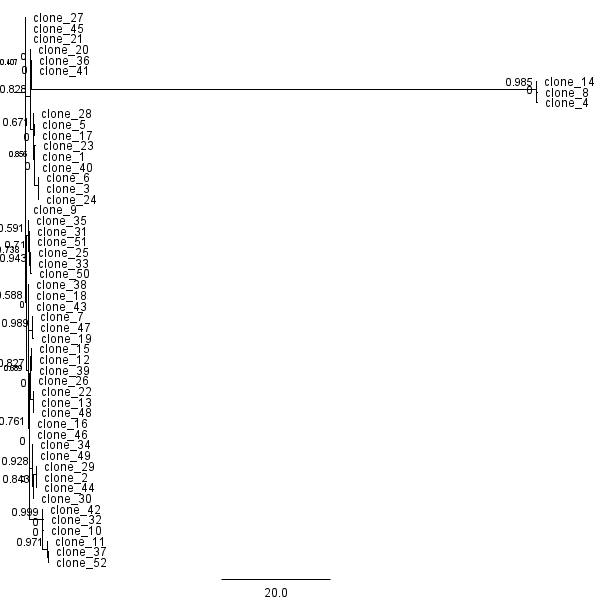

### Fig4d2.png

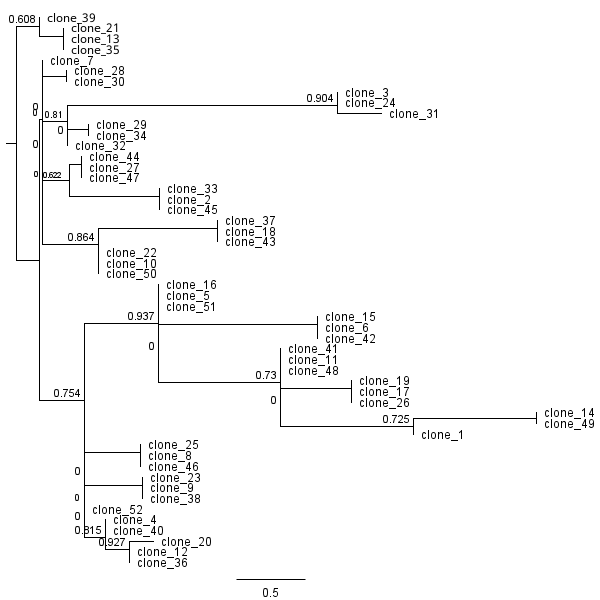

### Fig4d3.png

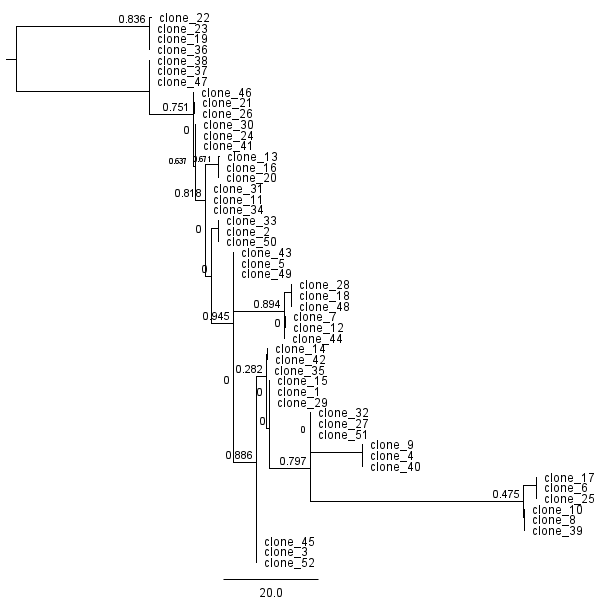

### Fig4d4.png

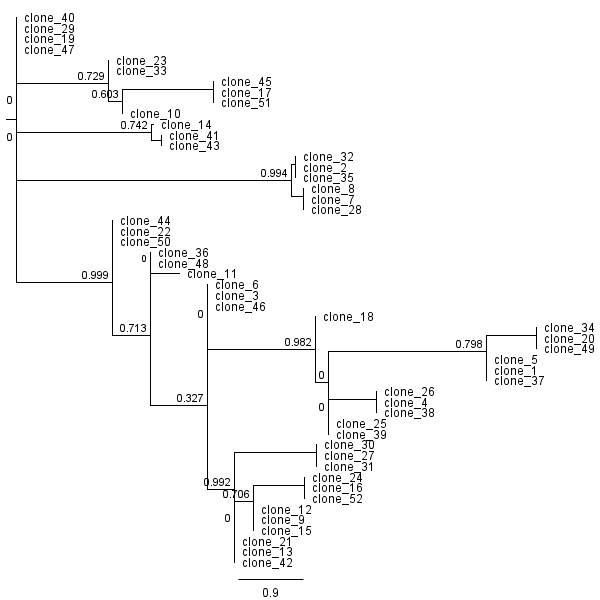

### Fig4d5.png

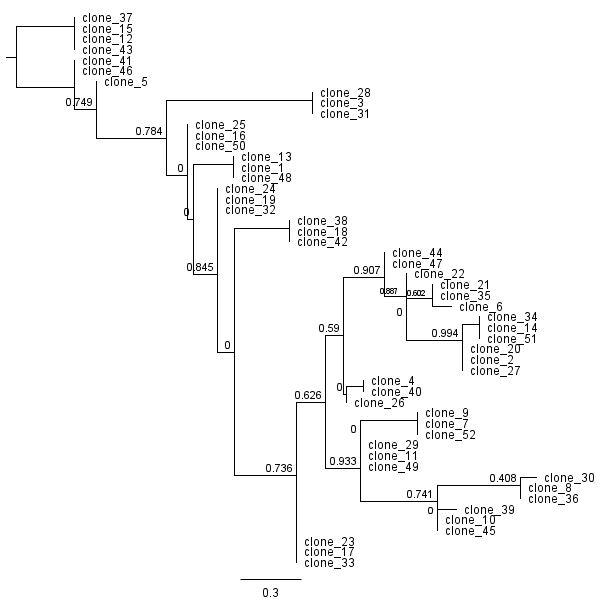

### Fig4e1.png

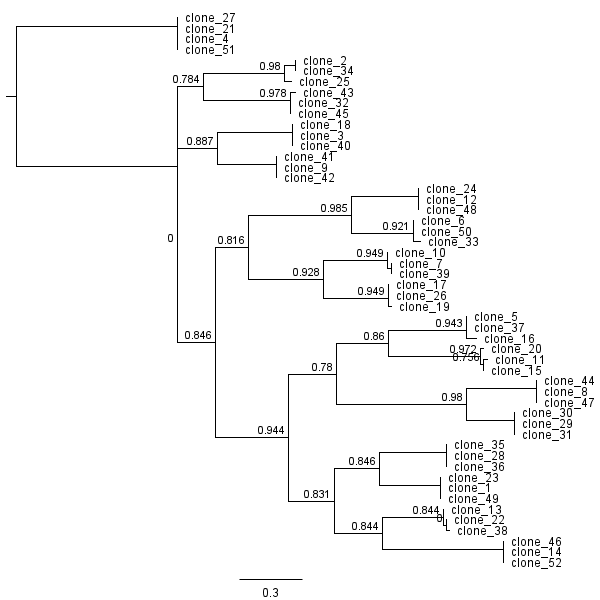

### Fig4e2.png

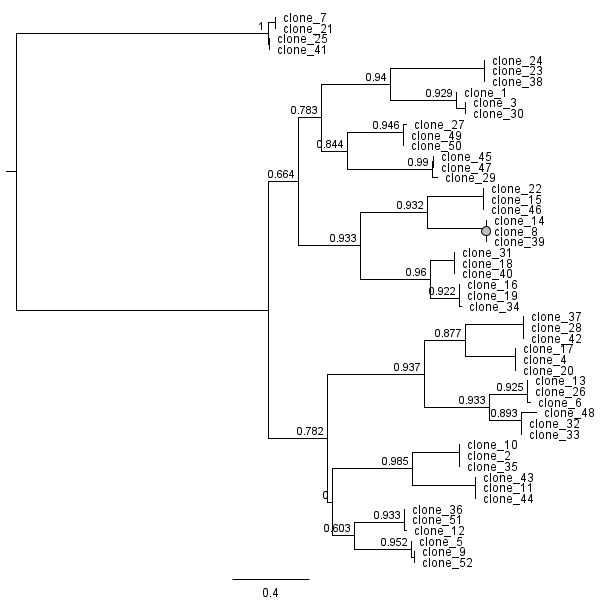

### Fig4e3.png

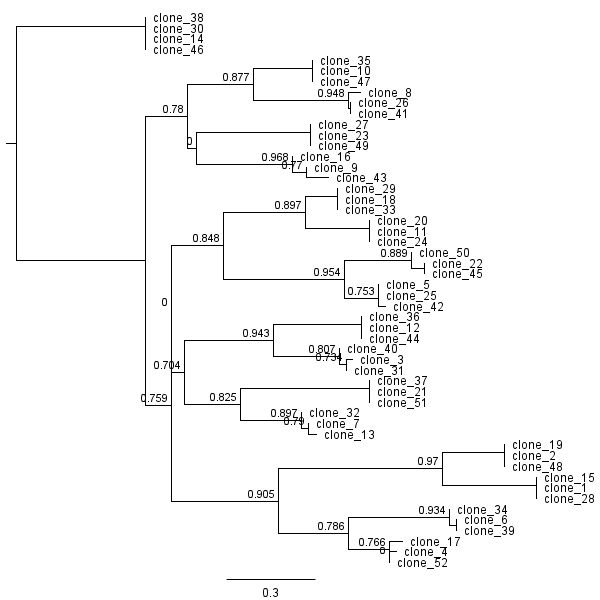

### Fig4e4.png

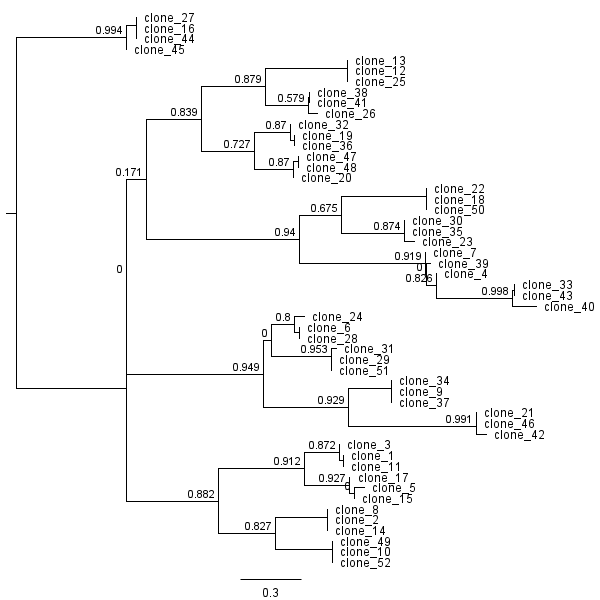

### Fig4e5.png

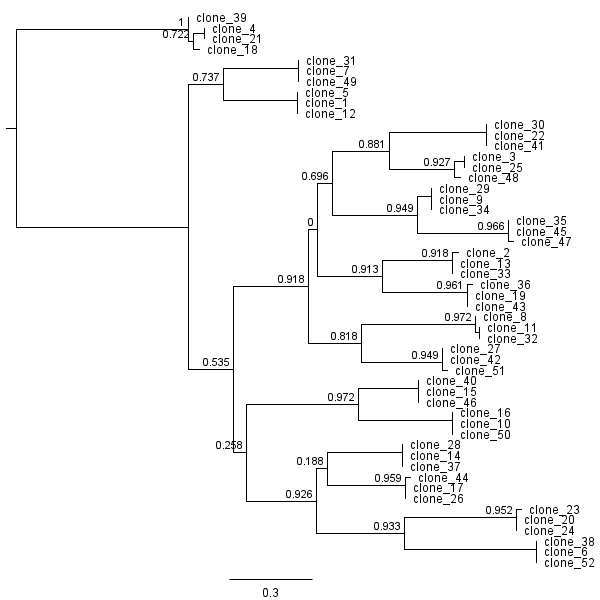
